# FOXO/DAF-16 modulates the transcription factor ROR/NHR-23 and inhibits the *let-7* microRNA to maintain multipotency during dauer

**DOI:** 10.64898/2026.07.18.739353

**Authors:** Himani Galagali, Matthew J. Wirick, Amelia F. Alessi, Margaret R. Starostik, Priya Balamurugan, Isabella Giudicelli Sims, Laurianne Pene, Ruhi Patel, Suhua Feng, Alison R. Frand, Steven E. Jacobsen, Xantha Karp, John K. Kim

## Abstract

Animals rapidly reprogram gene expression to adapt development to environmental stress. How gene regulatory programs that drive continuous development are repressed during stress-induced developmental arrest remains poorly understood. In *Caenorhabditis elegans*, starvation and overcrowding trigger entry into the stress-resistant, quiescent dauer stage. Here, we identify interactions among the conserved transcription factors DAF-16/FOXO and NHR-23/ROR, and the *let-7* family of microRNAs as key regulators of the switch from continuous development to dauer. We show that loss of *daf-16* during dauer causes elevated *let-7* family microRNAs and premature expression of the adult collagen reporter *col-19p*::GFP. Reducing *let-7* family activity suppresses this phenotype, whereas dauer-specific *let-7* expression is sufficient to induce *col-19p*::GFP expression. Mechanistically, DAF-16 inhibits *let-7* transcription in part by repressing *nhr-23*, which encodes a transcriptional activator of the *let-7* family and molting-cycle genes. ChIP-seq analysis reveals DAF-16 binding upstream of *nhr-23*, and *daf-16*; *daf-7* mutant dauers exhibit increased *nhr-23* mRNA and NHR-23 protein, supporting a model in which DAF-16 directly represses *nhr-23*. Integrated ChIP-seq and transcriptomic analyses identify 1,183 genes activated and 681 genes repressed by DAF-16 during dauer. Repressed targets are enriched for pro-growth genes involved in mitotic DNA replication and translational elongation. DAF-16 targets include 59 transcription factors that may mediate broader transcriptional reprogramming during dauer to maintain multipotency and establish quiescence. Together, these findings reveal that DAF-16/FOXO establishes stress-induced developmental arrest by coupling activation of protective pathways with repression of conserved developmental timing, growth, and differentiation programs.

**Significance statement:** Animals often pause development during environmental stress and then resume normal development when conditions improve. How developmental programs are temporarily halted without disrupting later cell fates remains poorly understood. We show that, during stress-induced dauer arrest in *C. elegans*, the conserved FOXO transcription factor DAF-16 represses the differentiation-promoting *let-7* microRNA pathway. DAF-16 inhibits the ROR homolog NHR-23, which normally activates *let-7* and molting-cycle genes. Genome-wide analyses further show that DAF-16 both activates stress-response genes and represses growth and developmental genes. These findings reveal how a conserved stress-responsive factor coordinates survival, developmental arrest, and maintenance of cellular multipotency.

## Introduction

The spatiotemporal gene expression programs that orchestrate animal development must adapt to fluctuating environmental conditions, such as changes in temperature or nutrient availability. For example, the nematode *Caenorhabditis elegans* (Cassada and Russell, 1975), the annual killifish *Nothobranchius furzeri* (Platzer and Englert, 2016), and several insects of the order Lepidoptera (reviewed in Denlinger, 2022, Schiesari and O’Connor, 2013) can enter a quiescent, stress-resistant state of developmental diapause during adverse environmental conditions. The signaling pathways that transduce environmental cues to control the decision to enter diapause are well studied (Riddle et al., 1981; Riddle and Albert, 1997; Fielenbach and Antebi, 2008). In contrast, the molecular mechanisms that redirect gene expression programs from continuous development to stress response remain poorly understood.

Entry into diapause causes developmental arrest or slows developmental progression (reviewed in Denlinger, 2022, Schiesari and O’Connor, 2013). Diapause thus interrupts ongoing developmental gene programs until organisms encounter favorable conditions. Remarkably, adult *C. elegans* that develop post-diapause exhibit no developmental defects and, in some cases, have enhanced fitness (Hall and Sengupta, 2010). To ensure developmental fidelity after diapause, entry into diapause must be coupled with mechanisms that maintain multipotent cell fates, sometimes for significant periods of time (Euling and Ambros, 1996; Liu and Ambros, 1991; Karp and Greenwald, 2013).

Under favorable conditions, *C. elegans* undergoes continuous development through four larval stages (L1–L4) before adulthood. Under unfavorable conditions such as starvation or overcrowding, juveniles enter an alternative quiescent, stress-resistant life stage called dauer. Here we investigate molecular mechanisms contributing to quiescence establishment, cell fate maintenance, and stress response during dauer.

During continuous development, the transition between larval stages is marked by a molt. Larvae molt at regular 8–10-hour intervals, as governed by the molting cycle timer. The PERIOD homolog LIN-42 functions as a component of the molting cycle timer (Monsalve et al., 2011). We previously reported that negative feedback between the Retinoid-related Orphan Receptor homolog NHR-23 and the microRNA *let-7* paces the molting cycle (Patel, Galagali et al., 2022). NHR-23 promotes *let-7* transcription, while *let-7* inhibits *nhr-23* translation by targeting its mRNA for degradation. Several molting genes, including *lin-42*, are transcriptionally activated by NHR-23 and post transcriptionally repressed by *let-7,* establishing oscillatory gene expression patterns. More than 3,700 genes, including 257 molting-associated genes, exhibit oscillatory expression during continuous development (Hendriks et al., 2014; Kim et al., 2013; Meeuse et al., 2020). Entry into dauer interrupts this rhythmic gene expression and transiently halts the molting cycle.

The heterochronic gene regulatory network, including the *let-7* family of microRNAs and their targets, controls another important aspect of *C. elegans* development: the timing of cell divisions and cell fate transitions in hypodermal seam cells (Ambros and Horvitz, 1984; Rougvie and Slack, 2001; Ambros and Ruvkun, 2018). During continuous development, stem cell-like progenitor cells in the hypodermis undergo specific patterns of symmetric and asymmetric divisions that renew stem cell populations while generating differentiated daughter cells (Sulston and Horvitz, 1976). The *let-7* paralogs *mir-48, mir-84 and mir-241* downregulate the mRNA encoding the transcription factor, HBL-1, to drive changes in seam cell fate as worms the transition from the L2 to L3 stage (Abbott et al., 2005). During the L4-to-adult transition, *let-7* degrades *lin-41* mRNA, which encodes an RNA binding protein. Repression of *lin-41* by *let-7* permits expression of the transcription factor LIN-29, which then activates adult collagens and specifies adult cell fates (Figure 1A; Ambros, 1989; Rougvie and Ambros, 1995; Reinhart et al., 2000). Entry into the dauer stage results in delay of the stereotyped pattern of hypodermal seam cell division and differentiation until worms encounter conditions that are favorable for development.

**Figure 1.**
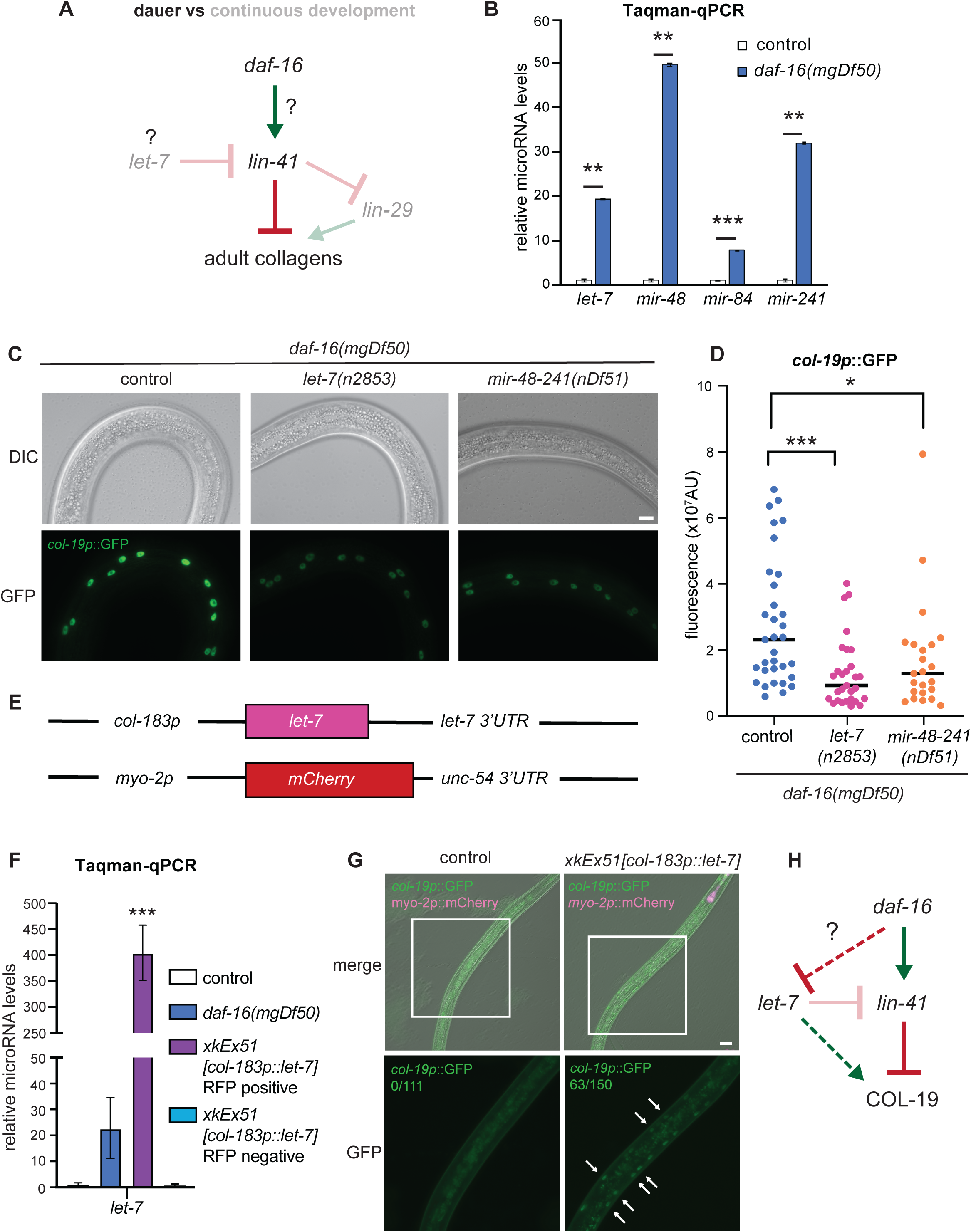
Upregulation of *let-7* family members drives precocious *col-19p*::GFP expression during dauer. **A.** Schematic representing the pathway that suppresses adult fates in dauer, as characterized in Wirick et al., 2021. Genetic interactions between *let-7, lin-41*, and *lin-29* that establish adult fates during continuous development are indicated with fainter text and arrows. All strains used in B-G have *daf-7(e1372); maIs105[col-19p::gfp]* in the background. **B.** Taqman-qPCR measuring levels of *let-7*, *mir-48*, *mir-84*, and *mir-241* microRNAs in control vs *daf-16(mgDf50)* dauers. Expression of each microRNA was normalized to U18 and then to the value in control dauers. Bars represent the mean of three biological replicates and error bars represent standard deviation. P values were calculated using Student’s t-test. ** p<0.01, *** p<0.001. **C.** Representative images of dauers of the indicated strains. Scale bar = 10 μm. **D.** Quantification of *col-19p*::GFP expression shown in C. Each data point represents the average fluorescence intensity from at least three hypodermal cells in individual dauers. Bars indicate the median of the distribution. N = 23–35. P values were calculated using the Mann-Whitney test. * p<0.05. *** p<0.001. **E.** Schematic of constructs used to overexpress *let-7* during dauer in *daf-7(e1372); maIs105[col-19p::gfp]; xkEx51[col-183p::let-7+myo-2p::mCherry]* animals. **F.** Taqman qPCR measuring *let-7* levels during dauer in the indicated genotypes. Expression was normalized to U18 and then to the value in control *dauers*. Bars represent mean of two technical replicates and error bars represent standard deviation. P values were calculated using Student’s t-test. *** p<0.001. **G.** Representative images of dauers of the indicated genotypes. The number of animals expressing *col-19p*::GFP is indicated. Scale bar = 20 μm. Images in the lower row show magnified regions highlighted by white boxes. Arrows indicate *col-19p*::GFP expression in hypodermal cells. **H.** Schematic summarizing the genetic interactions established in this figure. Dashed lines indicate possible involvement of additional factors. The question mark indicates the pathway investigated in Figure 2.

We previously reported that the FOXO homolog *daf-16* prevents precocious adult fates in hypodermal and seam cells during dauer by promoting *lin-41* expression (Wirick et al., 2021). DAF-16 is the conserved transcription factor downstream of the insulin signaling pathway. Under favorable conditions, insulin-like neuropeptides secreted by neurons bind to the receptor DAF-2 in multiple tissues, triggering a phosphorylation cascade that results in DAF-16 phosphorylation and its retention in the cytoplasm. In contrast, under unfavorable conditions, unphosphorylated DAF-16 and translocates to the nucleus and activates genes promoting stress response and dauer formation (reviewed in Murphy and Hu, 2013). Our previous findings (Wirick et al., 2021) suggested that *daf-16* plays a novel role in regulating heterochronic genes during dauer, beyond its established function in stress response.

Dauer larvae with loss-of-function alleles of *daf-16* precociously express several adult collagen genes, including *col-19*. Using a transgene expressing *col-19p::*GFP as an adult fate marker, we previously showed that depletion of *lin-41* also causes precocious adult fates (Figure 1A; Wirick et al., 2021).

In this study, we demonstrate that *daf-16* inhibits the *let-7* family of microRNAs to maintain hypodermal seam cell multipotency during dauer. This regulation occurs at the transcriptional level through the transcription factor NHR-23. ChIP-seq analysis reveals DAF-16 binding at the *nhr-23* promoter during dauer, consistent with direct transcriptional repression. Reduced *nhr-23* expression, in turn, prevents *let-7* transcription, thereby blocking premature adult differentiation programs. Genome-wide analysis identifies 1,183 genes activated and 681 genes repressed by DAF-16, with repressed targets enriched for pro-growth functions. These findings support a model in which DAF-16 maintains multipotency and enforces developmental quiescence by repressing the NHR-23/*let-7* developmental-timing pathway while coordinately activating stress-response programs.

## Results

### *let-7* family microRNAs are upregulated in *daf-16(-); daf-7(-)* dauers

During the L4-to-adult transition, *let-7* downregulates its target *lin-41* (Reinhart et al., 2000). Because 3ʹ UTR-mediated upregulation of *lin-41* partially suppressed precocious *col-19p*::GFP expression in *daf-16(-)* dauers (Wirick et al., 2021), we investigated whether the *let-7* family of microRNAs regulates multipotency during dauer (Figure 1A).

Larvae with loss-of-function alleles of *daf-16* do not enter dauer even in dauer-inducing environmental conditions. Therefore, we used the temperature sensitive dauer-constitutive allele *daf-7(e1372)* to study dauer larvae with the null allele *daf-16(mgDf50)*, hereafter referred to as *daf-7(-)* and *daf-16(-)*. Taqman qPCR showed that mature *let-7, mir-48, mir-84,* and *mir-241* were all significantly upregulated in *daf-16(-)*; *daf-7(-)* compared to *daf-7(-)* dauers at 52 h at 24°C (Figure 1B). Levels of *let-7* increased ∼20-fold, *mir-48* ∼50-fold, *mir-84* ∼8-fold, and *mir-241* ∼30-fold in *daf-16(-)*; *daf-7(-)* dauers compared to *daf-7(-)* dauers. These data suggest that *daf-16* inhibits accumulation of the *let-7* family during dauer.

To test whether this upregulation of the *let-7* family mediates precocious *col-19p*::GFP expression, we introduced the hypomorphic allele *let-7(n2853)* into *daf-16(-); daf-7(-)* dauers, resulting in a ∼2.6-fold decrease in *col-19p*::GFP expression (Figure 1C, D). Similarly, introduction of the loss-of-function allele *mir-48–241(nDf51)* in *daf-16(-); daf-7(-)* dauers led to a ∼1.8-fold decrease (Figure 1C, D). Because all *let-7* family members share the same seed sequence, redundancy in target silencing likely explains the incomplete suppression of the *col-19p*::GFP expression observed. Together, these data suggest that misexpression of the *let-7* family microRNAs is necessary for precocious *col-19p*::GFP expression in *daf-16(-); daf-7(-)* dauers.

We next tested whether upregulation of *let-7* is sufficient to induce precocious *col-19p*::GFP expression in dauers. In *C. elegans*, there are 2 primary transcripts for *let-7*, originating from different transcriptional start sites (Kai et al., 2013). A plasmid with the shorter primary *let-7* cloned downstream of the dauer-specific hypodermal promoter *col-183p* (Shih et al., 2019) was injected into *daf-7(-); col-19p::gfp* worms (Figure 1E–F, Figure S1A). Among *daf-7(-); col-19p::gfp* dauers expressing the *myo-2p*::RFP co-injection marker in the pharynx, 42% also expressed *col-19p*::GFP in hypodermal cells (Figure 1G). However, the intensity of *col-19p*::GFP in *daf-7(-); col-183p::let-7* dauers was lower than that observed in *daf-16(-); daf-7(-)* dauers (Figure 4B). These results indicate that overexpression of *let-7* is sufficient to induce *col-19p*::GFP expression during dauer. In summary, *let-7* family microRNAs are upregulated in *daf-16(-); daf-7(-)* dauers, and this increase is, in part, necessary and sufficient to induce precocious *col-19p*::GFP expression. These findings support a model in which *daf-16*-mediated inhibition of the *let-7* family suppresses adult differentiation programs and maintains seam cell multipotency during dauer (Figure 1H).

### *daf-16* inhibits *let-7* transcription

MicroRNAs are transcribed from endogenous loci as long primary transcripts that are subsequently processed by the Microprocessor complex and Dicer into precursor hairpins and mature microRNA duplexes, respectively. One strand of the mature duplex is then unwound and loaded into the Argonaute protein to form the core of the microRNA-induced silencing complex (miRISC). Each step in this biogenesis pathway is subject to regulation, which can lead to differential microRNA expression (reviewed in Ambros and Ruvkun, 2018; Ha and Kim, 2014).

Because *daf-16* inhibits accumulation of mature *let-*7 family microRNAs during dauer, we asked whether *daf-16* regulates *let-7* transcription. To test this, we used a strain expressing GFP under the *let-7* promoter (*let-7p::GFP*) in *daf-7(-)* and *daf-16(-)*; *daf-7(-)* backgrounds. GFP levels were measured from the L2d stage, preceding dauer entry, through the dauer stage. Beginning at 31 hours, *let-7p*::GFP expression was significantly higher levels in *daf-16(-)*;*daf-7(-)* animals than in *daf-7(-)* animals, resulting in an approximately 2-fold increase in GFP signal at the 48 hour time point when the animals are in the dauer stage (Figure 2A, B). These results are consistent with *daf-16* inhibiting *let-7* transcription during L2d.

**Figure 2.**
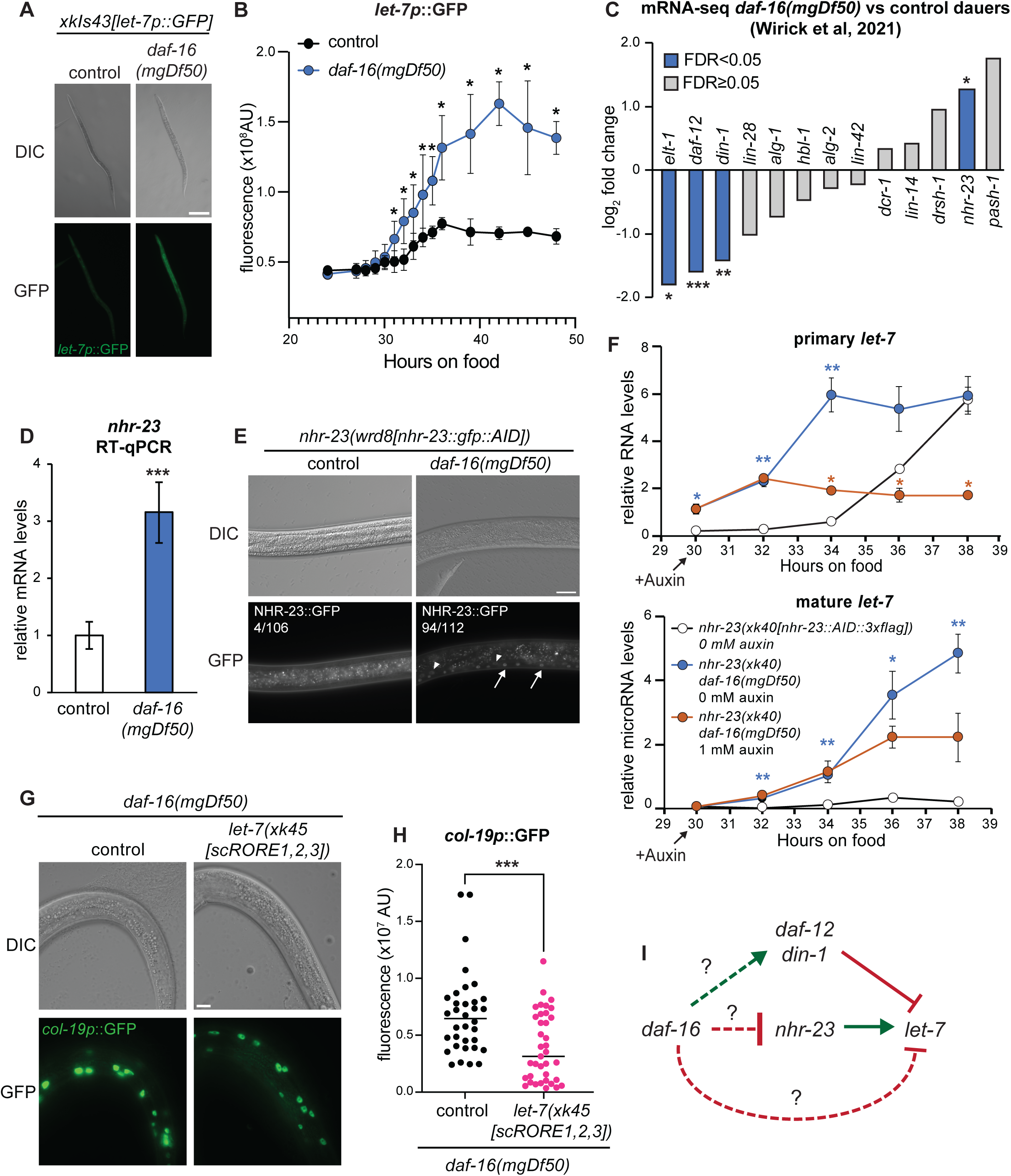
NHR-23 mediates upregulation of *let-7* family transcription in *daf-16(-); daf-7(-)* animals. All strains used in this figure have *daf-7(e1372)* in the background. **A.** Representative images of *xkIs43[let-7p::GFP]* and *daf-16(mgDf50); xkIs43[let-7p::GFP]* dauers at 48 h at 24°C. Scale bar = 100 μm **B.** Quantification of *let-7p::GFP* signal from 28–48 h during the L2d and dauer stages. Markers represent median measurements and error bars represent standard deviation. N = 15–25. P values were calculated by the Mann-Whitney test. *p<0.05. **C.** Differential gene expression analysis of previously characterized regulators of *let-7* microRNAs in *daf-16(mgDf50)* versus control dauers (Wirick et al., 2021). Blue bars data points indicate genes with average log_2_ fold change ≥ 1 and FDR ≤ 0.05 from 2 biological replicates. Grey bars indicate genes that did not meet these criteria. *FDR<0.05; **FDR<0.01; ***FDR<0.001. **D.** RT-qPCR measuring *nhr-23* mRNA levels in control versus *daf-16(mgDf50)* dauers. Expression values for were normalized to *eft-2* and then to the value in control dauers. Bars represent the mean of three biological replicates and error bars represent standard deviation. P values were calculated using Student’s t test. ***p<0.001; n.s., not significant. **E.** Representative images showing NHR-23::GFP expression in the indicated genotypes. The number of animals expressing NHR-23::GFP::AID is indicated. Arrows indicate hypodermal cells and arrowheads indicate seam cells. Scale bar = 20μm **F.** RT-qPCR measuring primary (top) and mature (bottom) *let-7* levels in the indicated genotypes. Strains have *ieSi57[eft-3p::tir1].* Worms were exposed to 0 mM or 1 mM auxin at 30 h. Each data point represents the mean of two technical replicates and error bars represent standard deviation. Blue asterisks indicate comparisons between *nhr-23::AID; tir1* 0 mM auxin and *nhr-23::AID daf-16(mgDf50)* 0 mM auxin. Orange asterisks indicate comparisons between *nhr-23::AID daf-16(mgDf50),* 0 mM auxin and *nhr-23::AID daf-16(mgDf50),* 1 mM auxin. P values were calculated using Student’s t test. *p<0.05, **p<0.01. **G.** Representative images of *col-19p*::GFP expression in the indicated genotypes with *maIs105[col-19p::gfp]*. Scale bar = 10 μm **H.** Quantification of *col-19p*::GFP expression shown in G. Each data point represents the average fluorescence intensity from at least three hypodermal cells in individual dauers. Bars represent the mean of the distribution. N = 33–35. P values were calculated using the Mann-Whitney test. ***p<0.001. **I.** Schematic summarizing genetic interactions established in this figure. Dashed lines indicate possible involvement of additional factors. The question mark indicates the pathway investigated in Figure 3.

Several factors regulate *let-7* transcription during continuous development. The transcription factors HBL-1 (Roush and Slack, 2009), LIN-14 (Tsialikas et al., 2017), and LIN-42 (McCulloch and Rougvie, 2014; Perales et al., 2014; Van Wynsberghe et al., 2014) inhibit *let-7* transcription, whereas ELT-1 (Cohen et al., 2015) and NHR-23 (Patel, Galagali et al., 2022) activate *let-7* transcription. Under favorable conditions, the nuclear hormone receptor DAF-12 binds its ligand and activates *let-7* expression. Under unfavorable conditions, however, DAF-12 interacts with its corepressor DIN-1 to inhibit *let-7* expression (Bethke et al., 2009; Hammell et al., 2009). To determine whether any of these regulators are affected by loss of *daf-16*, we examined our previously published mRNA-seq dataset comparing gene expression in *daf-16(-); daf-7(-)* and *daf-7(-)* dauers (Wirick et al., 2021). We focused on known regulators of *let-7* transcription and microRNA biogenesis factors rather than performing an unbiased screen in this analysis (Figure 2C). According to these data, *nhr-23* was significantly upregulated in *daf-16(-); daf-7(-)* dauers, whereas *daf-12* and *din-1* were significantly downregulated (Figure 2C). The transcriptional activator *elt-1* was also downregulated and was therefore excluded from further analysis. None of the other transcription factors or components of the microRNA biogenesis machinery were differentially expressed (Figure 2C, Figure S2A). The ∼3-fold upregulation of *nhr-23* and ∼2.5-fold downregulation of *daf-12* in *daf-16(-); daf-7(-)* dauers were validated by RT-qPCR (Figure 2D, Figure S2B). However, the apparent reduction in *din-1* expression was not statistically significant by RT-qPCR. Together, these data suggest that increased expression of the *let-7* transcriptional activator *nhr-23*, along with reduced transcript expression encoding the inhibitory DAF-12/DIN-1 heterodimer, may contribute to the upregulation of *let-7* transcription in *daf-16(-); daf-7(-)* dauers.

A strain expressing an endogenously tagged *nhr-23* allele, *nhr-23::gfp::flag::aid* (Auxin Inducible Degron) (Ragle et al., 2020, a gift from Jordan Ward), was used to determine if NHR-23 protein levels were elevated in *daf-16(-); daf-7(-)* dauers. All strains described below carry the *eft-3p::tir1* transgene, which enables degradation of NHR-23::GFP::FLAG::AID in somatic tissues upon auxin exposure. While only gut autofluorescence was detectable in the majority of *nhr-23::gfp::flag::aid; daf-7(-)* dauers, NHR-23::GFP::FLAG::AID was expressed in hypodermal and seam cell nuclei in 84% (94/112) of *nhr-23::gfp::flag::aid daf-16(-); daf-7(-)* dauers (Figure 2E). Exposure to 1mM auxin for 2 hours rendered the NHR-23::GFP::FLAG::AID signal undetectable in nearly all *nhr-23::gfp::flag::aid daf-16(-); daf-7(-)* dauers (Figure S2C).

These data indicate that in *daf-16(-); daf-7(-) dauers*, both *let-7* transcription and expression of the *let-7* transcriptional activator NHR-23 are upregulated, while components of the *let-7* transcriptional inhibitory complex, *daf-12* and *din-1*, are downregulated. These findings support a model in which, during dauer, *daf-16* represses *nhr-23* and may promote DAF-12/DIN-1-mediated inhibition of *let-7* transcription. However, a direct role for *daf-16* in repressing *let-7* transcription cannot be ruled out (Figure S2D).

### NHR-23 mediates upregulation of *let-7* transcription in *daf-16(-); daf-7(-)* dauers

The role of DAF-12 and DIN-1 in regulating *let-7* expression during the L2d stage has been characterized previously (Bethke et al., 2009; Hammell et al., 2009). NHR-23 has been identified as a transcriptional activator of *let-7* during continuous development (Kai et al., 2013; Patel et al., 2022). We next asked whether the upregulation of NHR-23 in *daf-16(-); daf-7(-)* animals mediates the increase in *let-7* transcription during L2d and dauer.

As *nhr-23* is an essential gene, we used CRISPR/Cas9-mediated genome editing to tag *nhr-23* at its endogenous locus with an *aid::3xflag* sequence. All strains discussed below also carry the *eft-3p::tir1* transgene, allowing conditional degradation of NHR-23::AID::3xFLAG in somatic tissues upon auxin treatment. We first confirmed that NHR-23::AID::3xFLAG can be conditionally depleted in dauers (Figure S3A).

To monitor primary and mature *let-7* levels*, nhr-23::AID::3xflag daf-16(-)*; *daf-7(-)* L2d animals were transferred to plates containing 0mM or 1mM auxin at 30 hours after plating embryos on food. Samples were collected every two hours for RNA extraction. As a control, a parallel set of *nhr-23::AID::3xflag; daf-7(-)* L2d animals maintained on 0mM auxin was collected. In the absence of auxin, primary *let-7* levels peaked earlier in *nhr-23::AID::3xflag daf-16(-); daf-7(-)* than in *nhr-23::AID::3xflag; daf-7(-)* controls (Figure 2F top, Figure S3B). In contrast, when NHR-23 was depleted by 1 mM auxin in *nhr-23::AID::3xflag daf-16(-); daf-7(-)* animals, primary *let-7* levels were consistently reduced compared to the 0mM auxin condition beginning at 34 hours (Figure 2F top, Figure S3B). Similarly, mature *let-7* accumulated at higher levels in *nhr-23::AID::3xflag daf-16(-); daf-7(-)* animals than in *nhr-23::AID::3xflag; daf-7(-)* controls. However, depletion of NHR-23 reduced mature *let-7* levels in *nhr-23::AID::3xflag daf-16(-); daf-7(-)* animals (Figure 2F bottom, Figure S3C). Similar trends were observed for mature *mir-48* (2/2 biological replicates), *mir-84* (2/2 biological replicates), and *mir-241* (1/2 biological replicates) (Figure S3D-E). Interestingly, despite the steady increase in primary *let-7* transcripts in *nhr-23::AID::3xflag; daf-7(-)* L2d larvae, mature *let-7* levels remained very low (Figure 2F, Figure S3B-C). This observation suggests that other *let-7* processing machinery may also be regulated by *daf-16* during dauer, potentially at the post-transcriptional level (Figure 2C).

We next used our *let-7p::GFP* reporter to monitor *let-7* transcription in *nhr-23::AID::3xflag; daf-7(-)* and *nhr-23::AID::3xflag daf-16(-); daf-7(-)* L2d animals with and without auxin. Depletion of NHR-23 in *nhr-23::AID::3xflag daf-16(-); daf-7(-),* starting at 30 hours resulted in a 1.5-fold reduction in *let-7p*::GFP levels at 38 hours (Figure S4A,B). However, both *nhr-23::AID::3xflag; daf-7(-)* and *nhr-23::AID::3xflag daf-16(-); daf-7(-)* animals exposed to auxin failed to form dauers, suggesting that some NHR-23 activity is required for dauer entry.

To further validate the role of *nhr-23* in regulating *let-7* levels in *daf-16(-)* dauers, we used an independent approach. *The let-*7 promoter contains three NHR-23 binding sites known as ROR binding elements (ROREs) (Figure S5A). Our previous work showed that scrambling two of these sites reduces NHR-23 binding at the *let-7* promoter and slows accumulation of mature *let-7* during continuous development (Patel, Galagali et al., 2022). Here, we scrambled all three ROREs using CRISPR/Cas9 genome editing to generate the *let-7(scRORE1,2,3)* allele. *let-7(scRORE1,2,3)* adults exhibited increased seam cell numbers, suggesting compromised *let-7* function (Figure S5B). RT-qPCR showed reduced amplitude of primary *let-7* expression, and Taqman-qPCR revealed slower accumulation of mature *let-7* during L3 development (Figure S5C-H). Mature *let-7* also accumulated more slowly in *daf-16(-); daf-7(-); let-7(scRORE1,2,3)* L2d animals (Figure S5I,J). Furthermore, consistent with reduced *let-7* activity, *daf-16(-); daf-7(-); let-7(scRORE1,2,3)* dauers exhibited approximately 2-fold lower levels of *col-19p*::GFP (Figure 2G,H), indicating that precocious *col-19p::*GFP expression requires adequate *let-7* expression, driven by NHR-23, during the L2d stage. In contrast, depletion of NHR-23 in *nhr-23::AID::3xflag daf-16(-); daf-7(-)* animals during the dauer stage itself did not affect *col-19p*::GFP expression (Figure S4C,D).

These data suggest that NHR-23-mediated activation of *let-7* transcription specifically during L2d drives precocious expression of *col-19p*::GFP in *daf-16(-); daf-7(-)* dauers. Overall, these findings support a model in which *daf-16* negatively regulates *nhr-23,* thereby inhibiting *let-7* expression and promoting seam cell multipotency (Figure 2I). However, the dauer-defective phenotype observed in *daf-7(-)* as well as *daf-16(-); daf-7(-)* animals upon depletion of NHR-23::AID::3xFLAG during L2d suggests that a basal level of NHR-23 is required for proper dauer entry. Thus, *daf-16* may not completely repress *nhr-23,* but instead modulates its expression levels to maintain multipotency of the hypodermal cells in dauer larvae.

### DAF-16 binds upstream of *nhr-23, daf-12, din-1,* and *let-7*

*daf-16* encodes a transcription factor. To investigate how *daf-16* regulates *nhr-23* expression, we performed chromatin immunoprecipitation followed by high-throughput sequencing (ChIP-seq) in starved dauers expressing *daf-16::mScarlet::3xflag* (a gift from Dr. Iva Greenwald, Figure S6H) and wild-type starved dauers, each in duplicate. To our knowledge, this dataset represents the first characterization of DAF-16 binding profiles during dauer. Consensus DAF-16 binding summits were identified as described in the Methods. These summits were located primarily in annotated promoter-Transcriptional Start Site(TSS) regions (58.4%) as well as intronic regions (16.8%) in both replicates (Figure S7C). As expected, strong DAF-16 binding summits were observed upstream of known targets *sod-3* and *mtl-1* (Figure S6A,B).

Two sets of DAF-16 binding summits were detected upstream of *nhr-23:* a stronger set within an intronic region, which is known to contain a TSS (Kostrouchova et al., 1998, Kostrouch et al., 1995), and a weaker set located further upstream (Figure 3A). Enrichment of DAF-16::3xFLAG at the intronic summits was confirmed by ChIP-qPCR (Figure S7A,B). A DAF-16 binding summit was also observed upstream of primary *let-7* (Figure 3B), although ChIP-qPCR indicated that the enrichment at the *let-7* promoter was lower than that observed at the *nhr-23* promoter (Figure S7A,B). Additional DAF-16 binding summits were also detected upstream of and within intronic regions of *din-1* (Figure S6C) and within intronic regions of *daf-12* (Figure S6D). DAF-16 binding was also found upstream of *mir-241* (Figure S6E) and *mir-84* (Figure S6F). As we previously reported (Wirick et al., 2021), DAF-16 binding summits were not detected upstream of *col-19* (Figure S6G).

**Figure 3.**
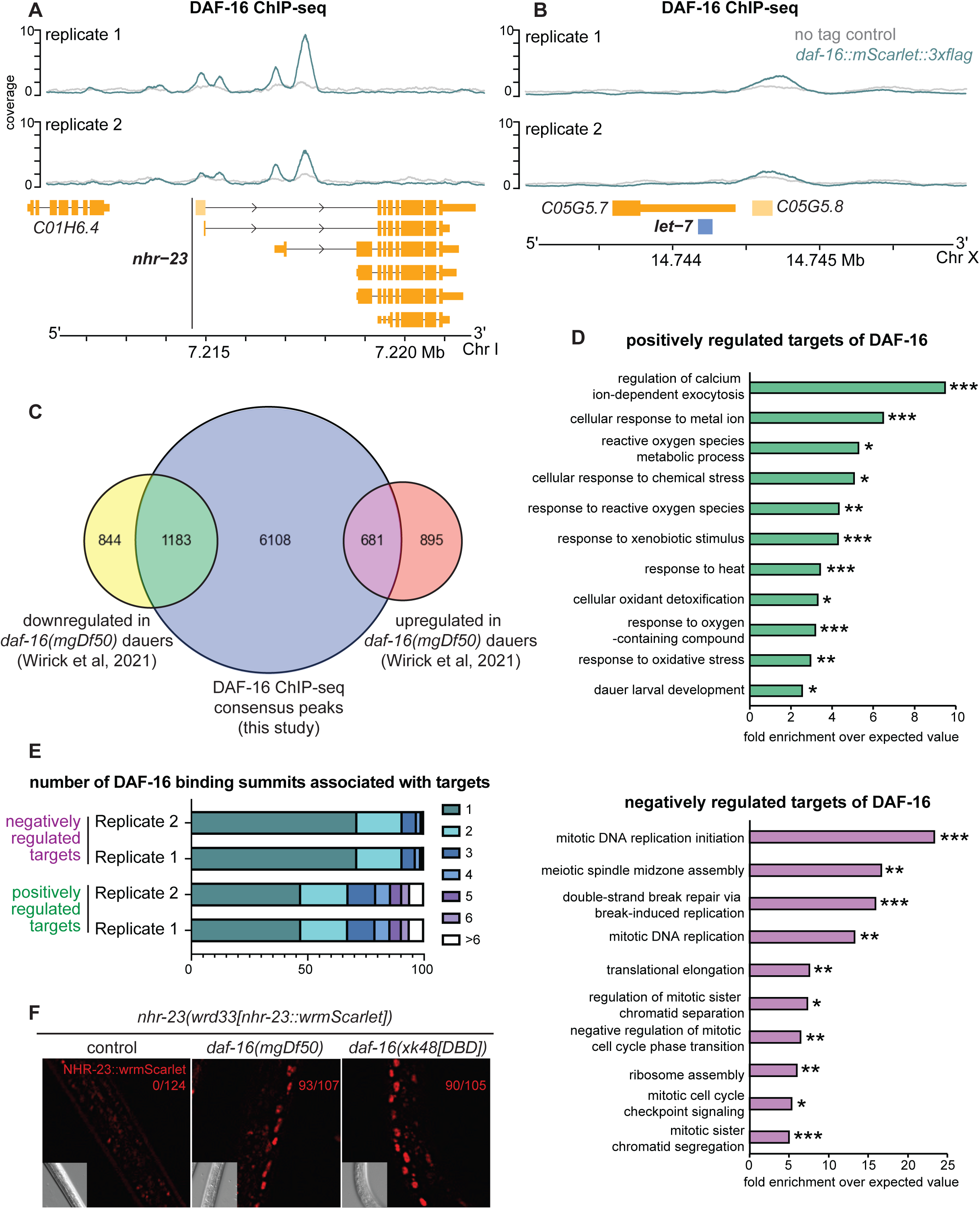
DAF-16 binds upstream of *nhr-23* and *let-7* during dauer. **A, B.** ChIP-seq tracks for N2 and *daf-16(ar620[daf-16::zf1-wrmScarlet-3xFLAG])* dauers showing DAF-16 binding summits upstream of **A.** *nhr-23* and **B.** *let-7*. **C.** Comparative analysis of predicted DAF-16 targets identified by ChIP-seq and differentially expressed genes from the mRNA-seq dataset published in Wirick et al., 2021. **D.** Gene ontology analysis of genes positively regulated by DAF-16 (top) and negatively regulated by DAF-16 (bottom). P values were calculated using Fisher’s exact test. *p<0.05, **p<0.01, ***p<0.001, ****p<0.0001. **E.** Distribution of the number of DAF-16 binding summits associated with positively or negatively regulated DAF-16 targets. F. Representative images of NHR-23::mScarlet expression in *daf-7(e1372)* induced dauer in the indicated genotypes. The number of dauers expressing NHR-23:: mScarlet is indicated.

These data indicate that DAF-16 binds upstream of specific isoforms of *nhr-23, daf-12* and *din-1*. While DAF-16 binding upstream of *daf-12* and *din-1* correlates with transcriptional activation, binding upstream of *nhr-23* correlates with transcriptional repression (Figure 2C). DAF-16 binding upstream of *let-7, mir-241* and *mir-84* may similarly contribute to transcriptional repression.

### DAF-16 plays a dual role as a transcriptional activator and suppressor

To identify biologically significant DAF-16 targets, we cross-referenced DAF-16 ChIP-seq dataset with our previously published mRNA-seq dataset comparing *daf-7(-)* and *daf-16(-); daf-7(-)* dauers (Wirick et al., 2021) (Figure 3C). Genes that have DAF-16 consensus binding summits in both dauer ChIP-seq replicates and that exhibit differential expression in *daf-16(-); daf-7(-)* versus *daf-7(-)* dauers were designated as DAF-16 targets. A total of 1183 DAF-16 targets, including *daf-12* and *din-1*, were downregulated in *daf-16(-); daf-7(-)* dauers and represent genes that are activated or positively regulated by DAF-16 (Dataset S1). In contrast, 681 DAF-16 targets, including *nhr-23*, were upregulated in *daf-16(-); daf-7(-)* dauers and represent genes that are inhibited or negatively regulated by DAF-16 (Dataset S2).

Gene ontology analysis revealed that the positively regulated targets of DAF-16 are enriched for genes involved in stress response (e.g., *mtl-1, sod-3, skn-1*) and dauer larval development (e.g., *sma-5, daf-12, daf-14*) (Figure 3D top, Dataset S3). In contrast, the negatively regulated targets are enriched for genes involved in DNA replication initiation (e.g., *mcm-2, mcm-3, mcm-4*), meiotic spindle assembly (e.g., *bub-1*), and translational elongation (e.g., *eef-1G, eef-1B.1*) (Figure 3D bottom, Dataset S4). These analyses suggest that, in addition to activating stress-response genes, DAF-16 also represses genes required for cell division and growth, consistent with a role for DAF-16 in pausing developmental gene expression programs in dauer.

Previous studies identified genes regulated by DAF-16 during aging (Murphy et al., 2003). Genes whose expression increased upon *daf-16* activation were designated Class I targets, whereas genes repressed by *daf-16* were classified as Class II targets. Notably, there is limited overlap between these aging-associated targets and the dauer-associated targets identified here (Figure S7D), suggesting that DAF-16 regulates distinct transcriptional programs during dauer and aging.

To investigate how DAF-16 might function both as a transcriptional activator and repressor, we examined features of DAF-16 binding at the two classes of targets. MEME-ChIP analysis identified a binding motif similar to the previously described DAF-16 associated element (DAE), CTTATCA (Tepper et al., 2013), among both positively and negatively regulated targets (Figure S7E). Intriguingly, we found differences in the number of DAF-16 binding summits associated with the two classes. While 71.3% of negatively regulated targets were associated with a single DAF-16 binding summit (487/683 for replicate 1 and 488/684 for replicate 2), 54.0% of positively regulated targets were associated with more than one DAF-16 binding summit (625/1186 for replicate 1 and 656/1185 for replicate 2) (Figure 3E). These observations suggest that the stoichiometry of DAF-16 binding at promoter elements may influence whether a target gene is activated or repressed.

Because DAF-16 has traditionally been characterized as a transcriptional activator, we sought to validate our findings using a novel DNA-binding-defective allele of *daf-16(xk48[daf-16::3xflag(DBD)])* (hereafter referred as *daf-16(DBD)*). This allele was generated by replacing four conserved residues (N215, R218, H219, S222) in the FOXO DNA-binding domain with alanine (Tsai et al., 2007) (Figure S8A) and includes a 3xFLAG tag for detection. Western blot analysis confirmed that DAF-16::3xFLAG(DBD) is expressed at levels comparable to wild-type DAF-16::3xFLAG (Figure S8B). Precocious *col-19p*::GFP expression during dauer was observed in *daf-16::3xflag(DBD); daf-7(-)* animals (Figure S8C), suggesting that *daf-16::3xflag(DBD*) behaves as a loss-of-function allele. Furthermore, using an *nhr-23::mScarlet* reporter allele (a gift from Jordan Ward), we found that NHR-23::mScarlet is inappropriately expressed in *daf-16::3xflag(DBD); daf-7(-)* dauers (Figure 3F). The presence of strong DAF-16 binding summits upstream of *nhr-23*, together with increased NHR-23 protein expression in two independent *daf-16* loss-of-function alleles, supports a model in which DAF-16 directly represses *nhr-23* expression during dauer.

### DAF-16-mediated repression of *nhr-23* limits expression of molting-cycle genes during dauer

DAF-16 targets include 59 genes annotated as transcription factors by gene ontology (Table S1). As DAF-16 enters the nucleus in response to unfavorable environmental conditions, regulation of these transcription factors by DAF-16 may help redirect gene expression programs from continuous development to dauer quiescence. To test this hypothesis, we focused on *nhr-23* because NHR-23 activates *let-7* transcription and molting-cycle genes during continuous development (Patel et al., 2022).

During continuous development, NHR-23 activates transcription of several genes that drive the molting cycle (Kostrouchova et al., 1998; Kouns et al., 2011). To test whether modulation of *nhr-23* by DAF-16 contributes to the switch between continuous development and dauer formation, we conditionally depleted NHR-23 from the soma during L2d in *nhr-23::AID::3xflag daf-16(-); daf-7(-)*animals, as described above, and performed RNA-seq. All strains used below contain *ieSi57[eft-3p::tir1]*.

We focused on NHR-23 targets previously characterized as Clock Controlled Genes (CCGs) (Patel et al., 2022) (Figure 4A). These genes share several defining features: 1) their expression oscillates with an 8–10 hour period during larval development (Hendriks et al., 2014; Kim et al., 2013); 2) they are associated with molting; 3) NHR-23 binds upstream of these genes based on modENCODE ChIP-seq dataset (Celniker et al., 2009; Gerstein et al., 2010); 4) the NHR-23 binding regions contain an enrichment of RORE motifs; and 5) knockdown of *nhr-23* by RNAi reduces their expression (Kouns et al., 2011). 46/54 CCGs were detected during L2d. Although these genes oscillate during continuous development, they displayed more variable expression patterns in L2d. This variability may reflect partial synchrony of the population, technical limitations in sampling resolution, the presence of both committed and uncommitted L2d animals in the sample, or physiological differences between L2d and continuously developing larvae. Three broad categories of gene expression were observed.

**Figure 4.**
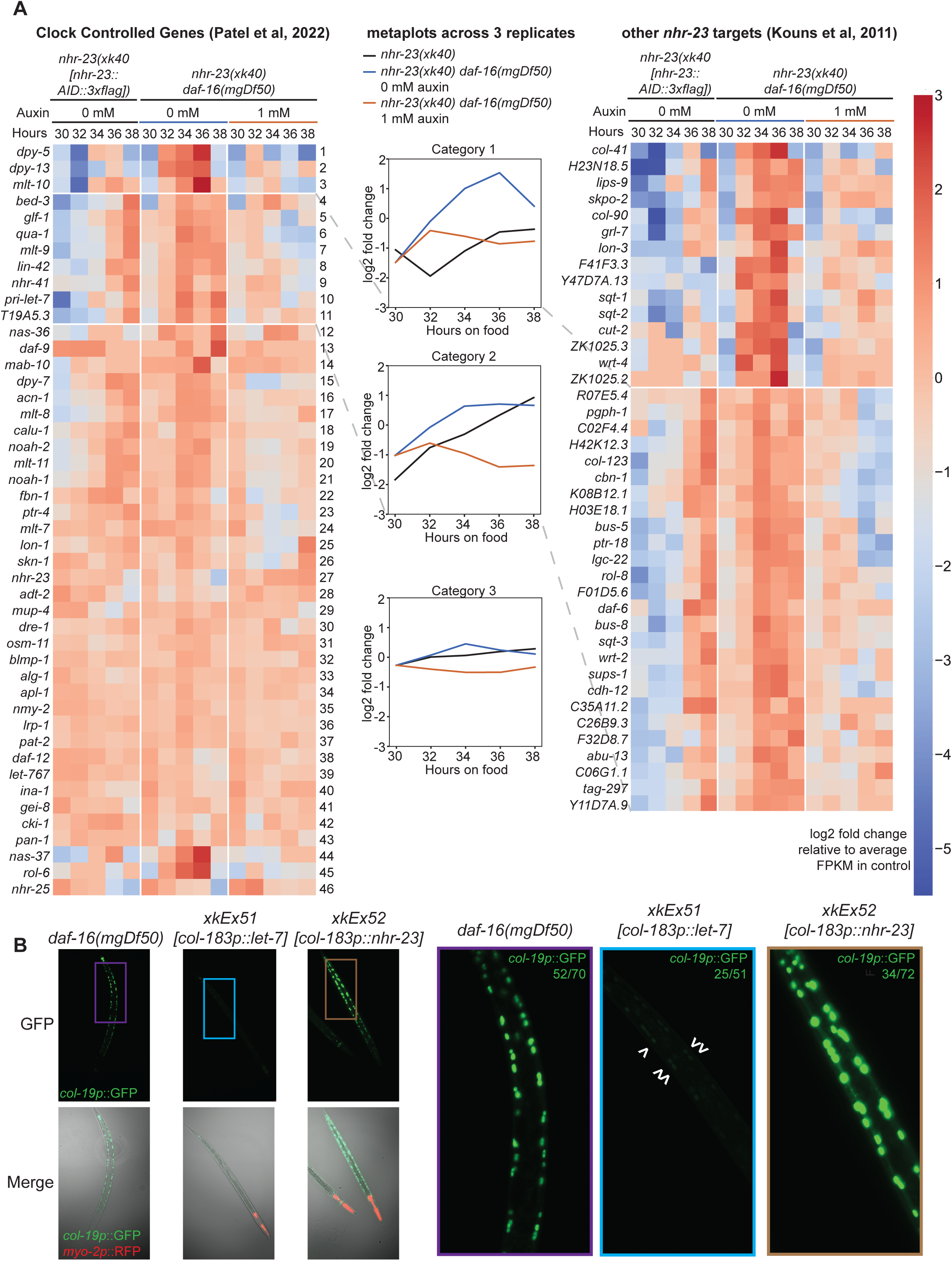
Suppression of *nhr-23* by DAF-16 prevents precocious adoption of adult fates and modulates expression of genes associated with continuous development. All strains have *daf-7(e1372)* in the background **A.** Heat maps showing expression profiles of *nhr-23* target genes classified as Clock Controlled Genes (CGC) (Patel et al., 2022) (left) and other *nhr-23* targets (Kouns et al., 2011) in the indicated genotypes during L2d development. Animals were transferred to plates containing 0 mM or 1 mM auxin at 30 h and maintained at 24°C. Metaplots represent average relative gene expression across all Category 1, 2, 3 genes across 3 biological replicates. **B.** Representative images of dauers of the indicated genotypes. Images on the right show magnified regions corresponding to the colored boxes. Arrowheads indicate faint *col-19p*::GFP expression in the hypodermal cells in *let-7*–overexpressing dauers.

Three CCGs *dpy-5*, *dpy-13* and *mlt-10* (rows1-3 in Figure 4A left, Figure S9A) represent Category 1 genes. Transcripts of *dpy-5*, *dpy-13* and *mlt-10* exhibited oscillatory expression patterns in *nhr-23::AID::3xflag; daf-7(-)* L2d animals. Expression of these genes in *nhr-23::AID::3xflag daf-16(-); daf-7(-)* animals was also oscillatory, but with increased amplitude. Auxin-mediated depletion of NHR-23 reduced the amplitude of expression of these genes, suggesting that elevated NHR-23 levels in *daf-16(-); daf-7(-)* animals promote their expression. A subset of previously described *nhr-23* targets (Kouns et al., 2011) also exhibited similar gene expression patterns (Figure 4A right). Notably, metaplots generated by averaging relative expression values of all Category 1 genes across 3 biological replicates suggest that depletion of NHR-23 in *daf-16(-)* L2ds restore the expression of these genes to the levels in the control strain (Figure 4A middle).

Seven CCGs, including *pri-let-7* and the PERIOD homolog *lin-42* (rows 4-11 Figure 4A left, Figure S9A), represent Category 2 genes. *lin-42* showed steadily increasing expression in both *nhr-23::AID::3xflag; daf-7(-)* and *nhr-23::AID::3xflag daf-16(-); daf-7(-)* L2d animals (row 8). However, *lin-42* expression reached higher levels earlier in *nhr-23::AID::3xflag daf-16(-); daf-7(-)* compared to *nhr-23::AID::3xflag; daf-7(-)* animals. Depletion of NHR-23 reduced the expression of these transcripts, suggesting that elevated NHR-23 levels in *daf-16(-); daf-7(-)* L2ds likely account for this earlier increase. Category 2 genes could also be identified among other *nhr-23* targets (Kouns et al., 2011) (Figure 4A right). In contrast to Category 1 genes, depletion of NHR-23 in *daf-16(-)* L2ds results in a sharp decrease of Category 2 targets compared to the control strain (Figure 4A middle).

Expression of Category 3 genes did not significantly change in developing *nhr-23::AID::3xflag; daf-7(-)* L2d animals (rows 12-46 Figure 4A left, Figure S9A). Some of these genes showed slightly elevated expression in *nhr-23::AID::3xflag daf-16(-); daf-7(-)* L2ds in an NHR-23-dependent manner (Figure 4A middle).

Together, these data suggest that misregulation of *nhr-23* in *daf-16(-); daf-7(-)* L2d animals and dauers leads to aberrant expression of genes normally associated with molting during continuous development. However, excepting transcripts in Category 1, depletion of NHR-23 in *daf-16(-); daf-7(-)* animals did not restore expression of these genes to levels observed in *daf-7(-)* dauers. This may support our previous observation that *daf-16(-); daf-7(-); nhr-23::AID::3xflag; eft-3p::tir1* animals are dauer defective on auxin and may therefore enter a developmental state that does not physiologically correspond to *daf-7(-)* dauers. Therefore, regulation of the key transcription factor *nhr-23* by DAF-16 during L2d and dauer may play a critical role in controlling levels of several molting cycle associated genes.

We next examined the physiological consequences of overexpressing *nhr-23* independent of *daf-16* during dauer. A construct driving *nhr-23* expression under the dauer-specific hypodermal promoter *col-183p* (Shih et al., 2019) was injected into *daf-7(-); col-19p::gfp* worms. Approximately 47% (34/72) of dauers overexpressing *nhr-23* also precociously expressed *col-19p*::GFP. Notably, the intensity of *col-19p*::GFP in *daf-7(-); col-183p::nhr-23* dauers was much higher than that observed in *daf-16(-); daf-7(-)* dauers (Figure 4B). Taqman qPCR revealed that expression of all *let-7* family members (*let-7, mir-48, mir-84* and *mir-241*) was elevated in *daf-7(-); col-183p::nhr-23* dauers (Figure S9B), which may partially explain the stronger *col-19p*::GFP signal. Together, these data indicate that overexpression of *nhr-23* is sufficient to induce precocious *col-19p*::GFP expression and suggest that modulation of *nhr-23* levels by DAF-16 plays an important role in limiting developmental progression programs during dauer. Similarly, DAF-16-mediated regulation of other transcription factors may contribute to rewiring of developmental gene programs during dauer.

## Discussion

We provide evidence that the conserved FOXO transcription factor DAF-16 inhibits transcription of the *let-7* family of microRNAs and other key developmental genes to maintain multipotency in hypodermal cells during dauer. DAF-16 binds upstream of the *let-7* family transcriptional regulators *nhr-23, daf-12* and *din-1* and modulates their expression, thereby suppressing accumulation of the *let-7* family during dauer. Repression of *nhr-23* by DAF-16 may also contribute to inhibition of molting-cycle components associated with continuous development or inappropriate molts from the dauer stage in unfavorable conditions. Together, these findings suggest that DAF-16 plays a dual role as both a transcriptional activator and repressor to coordinate stress-responsive gene expression while actively pausing developmental programs during dauer (Figure S10).

### DAF-16 inhibits *let-7* transcription to maintain multipotency during dauer

In *C. elegans, let-7* and its paralogs are part of the heterochronic gene regulatory network that coordinates a series of stage-specific cell fate transitions during larval development. Entry into dauer interrupts these developmental programs. While transcriptional and post-transcriptional regulators of *let-7* during continuous development have been characterized, the mechanisms controlling *let-7* levels during dauer have remained largely unknown.

Here, we show that *daf-16* inhibits accumulation of the *let-7* family during dauer (Figure 1B, Figure 2A-B). We further demonstrate that levels of the *let-7* transcriptional activator NHR-23 are elevated in *daf-16(-)* dauers (Figure 2C-E), while mRNA levels of the transcriptional repressor complex components *daf-12,* and possibly *din-1*, are reduced in *daf-16(-)* dauers (Figure 2C, Figure S2B). Increased NHR-23 expression in *daf-16(-)* dauers promotes transcription of *let-7* and accumulation of mature *let-7* family members (Figure 2F, Figure S3). During dauer, DAF-16-mediated reduction of *let-7* activity allows continued expression of its target, the RNA-binding protein LIN-41, thereby suppressing precocious adoption of adult cell fates.

The delayed accumulation of primary *let-7* transcripts, but not mature *let-7,* during L2d in *daf-7(-)* animals (Figure 2F) suggests that components of the *let-7* processing machinery may also be regulated by DAF-16. One possible regulator is LIN-28, which binds *pri-let-7* and prevents its processing during L1 and L2 stages. Although the mechanisms controlling *lin-28* expression remain unclear, the nucleocytoplasmic distribution of LIN-28 has been proposed to modulate its activity (Van Wynsberghe et al., 2011). While *lin-28* mRNA levels remain unchanged between *daf-16(-); daf-7(-)* vs *daf-7(-)* dauers (Figure 2C), it remains possible that DAF-16 influences LIN-28 localization or activity. If so, this would represent an additional mechanism through which DAF-16 could modulate *let-7* levels. Importantly, mature *let-7* accumulates in *daf-7(-); col-183p::let-7b* dauers (Figure 1F), suggesting that mechanisms preventing *let-7* processing operate primarily during the L2d stage. This observation highlights the continued importance of transcriptional regulation of *let-7* by DAF-16 during dauer.

The *let-7* family of microRNAs plays conserved roles in promoting cellular differentiation across the animal kingdom (Galagali and Kim, 2020; Pasquinelli et al., 2000; Roush and Slack, 2008). In contrast, FOXO proteins have been widely characterized as regulators of stem cell quiescence and plasticity in diverse organisms (Boehm et al., 2012; Liang and Ghaffari, 2018). Consistent with this role, DAF-16 suppresses precocious adoption of adult fates in vulval precursor cells (VPCs) during dauer by inhibiting EGFR/Ras and LIN-12/Notch signaling (Karp and Greenwald, 2013). Bioinformatic analyses have also suggested that FOXO may directly regulate *let-7* transcription in mammalian systems (Gaeta et al., 2017). Although we cannot rule out the possibility that DAF-16 directly represses *let-7* during dauer, the presence of multiple regulatory mechanisms, including repression of *nhr-23* and activation of *daf-12* and *din-1*, may contribute to the robustness of developmental programs (Cassidy et al., 2019). Thus, our findings provide molecular evidence that environmental stress can repress the conserved, differentiation-promoting *let-7* family through the conserved, quiescence-associated FOXO transcription factor.

### DAF-16-mediated modulation of *nhr-23* may pause the molting cycle during dauer

We recently described a self-sustaining genetic oscillator involving NHR-23 and *let-7* that regulates the molting cycle during continuous development (Patel, Galagali et al., 2022). Here, we show that DAF-16 binds upstream of *nhr-23* and inhibits its expression during dauer. DAF-16-mediated repression of *nhr-23* could therefore stop the activity of the NHR-23–*let-7* genetic oscillator. Several molting cycle-associated genes are transcriptional targets of NHR-23 and post-transcriptional targets of *let-7* (Patel, Galagali et al., 2022). We show that some of these genes, including the cuticle components *mlt-10* and *dpy-13* and the PERIOD homolog *lin-42,* are misexpressed in *daf-16(-)* dauers in an NHR-23-dependent manner. Thus, DAF-16-mediated suppression of the NHR-23–*let-7* genetic oscillator may help pause the molting program and maintain quiescence during dauer. However, the dauer-defective phenotype observed when NHR-23::AID is depleted in *daf-7(-)* dauers suggests that some NHR-23 activity is required for dauer entry. Residual NHR-23 may drive the expression of dauer-enriched collagens (Wirick et al., 2021) or genes associated with the dauer molt.

### DAF-16 plays a dual role as a transcriptional activator and repressor

We previously reported that seam cells in *daf-16(-)* dauers continue to divide and fail to enter quiescence (Wirick et al., 2021). Our current data suggest a possible mechanism for this finding, as DAF-16 binds upstream of and negatively regulates genes involved in DNA replication and cell division during dauer (Figure 3D). Elevated expression of these genes in *daf-16(-)* dauers may therefore contribute to the failure of seam cells to undergo arrest. Regulation of an additional 59 transcription factors by DAF-16 may similarly contribute to the switch between continuous development and diapause.

Previous mRNA-seq analyses comparing *daf-2(-)*, *daf-16(-); daf-2(-)*, and wild type adults identified genes that were upregulated in *daf-2(-)* animals in a *daf-16-*dependent manner (Class I genes) and genes that were downregulated (Class II genes) (Murphy et al., 2003). Subsequent ChIP-seq analyses revealed that DAF-16 binds upstream of Class I genes through the consensus DAF-16 binding element (DBE), T(G/A)TTTAC. In contrast, the complementary transcription factor PQM-1 binds upstream of the Class II genes through the DAF-16 associated element (DAE), CTTATCA (Tepper et al., 2013). In our dataset, we did not observe enrichment for the DBE motif upstream of positively regulated DAF-16 targets. Instead, the DAE motif was enriched upstream of both positively and negatively regulated DAF-16 targets. Furthermore, RNAi-mediated knockdown of *pqm-1* did not induce precocious *col-19p::*GFP expression during dauer (Figure S4E), suggesting that PQM-1 does not mediate suppression of adult fates in dauer.

One possible explanation is that DAF-16 associates with distinct cis-regulatory elements and trans-acting factors to function as both a transcriptional activator and repressor during dauer. Similar mechanisms have been described for other Forkhead domain proteins that possess both activating and repressing functions. For example, in regulatory T cells, FOXP3 functions as a transcriptional activator when associated with other transcription factors RELA, IKZF2 and KAT5, but acts as a transcriptional repressor when associated with the histone methyltrasferase EZH2 and transcription factors YY1 and IKZF3 (Kwon et al., 2017). Other Forkhead proteins FOXD3 and FOXP1 repress transcription by recruiting the corepressors Groucho and CTBP1, respectively (Li et al., 2004; Yaklichkin et al., 2007). Interactions with different cis-regulatory elements and trans-acting factors may also explain the differential stoichiometry of DAF-16 binding that we observed between positively and negatively regulated targets of DAF-16 (Figure 3E). Although the identity of these trans-acting factors remains unknown, DAF-16 has been shown to interact with HLH-30 under various stress conditions to modulate gene expression programs (Lin et al., 2015).

An alternative possibility is that nuclear DAF-16 inhibits transcriptional activators through protein-protein interactions rather than direct DNA binding. A similar mechanism has been described in smooth muscle cells, where FOXO4 interacts with the transcriptional activator myocardin to suppress differentiation. This effect is reversed by the addition of insulin-like growth factor and independent of the FOXO4 DNA binding domain (Liu et al., 2005). However, the robust DAF-16 binding observed upstream of repressed targets in our ChIP-seq dataset argues against this mechanism as the primary mode of repression during dauer.

In summary, DAF-16 promotes multipotency in dauer by modulating expression of *nhr-23*, *daf-12*, and *din-1* and by inhibiting the *let-7* family of microRNAs. Regulation of multiple transcription factors by DAF-16 may further contribute to the developmental switch from continuous growth programs to those associated with stress adaptation and dauer entry. Together, our findings suggest that stress-activated DAF-16 rewires developmental gene regulatory networks by repressing differentiation-promoting programs such as the NHR-23–*let-7* pathway, thereby preserving cellular multipotency and establishing developmental quiescence during dauer.

## Materials and Methods

### Working with *C. elegans* and isolation of dauers

*C. elegans* were cultivated, preserved, observed, and transformed using standard methods (Stiernagle, 2006). All strains used in this study are listed in Table S2. *xkIs43[let-7p::GFP]* was generated by UV-crosslinking and backcrossing a multicopy extrachromosomal strain gifted by Oliver Hobert.

Strains carrying *daf-7(e1372)* were maintained at 15°C and 20°C. Experiments examining *daf-7(e1372)* dauers were performed at 48-52 h at 24°C or 25°C (Fig 3F, Supp Fig 8C). For experiments comparing strains with *daf-7(e1372),* worms were developmentally semi-synchronized by sodium hypochlorite treatment. Briefly, eggs were isolated by lysing gravid hermaphrodites in sodium hypochlorite, suspended in M9 buffer, and plated on nematode growth medium (NGM) plates seeded with *E. coli* strain HB101 or OP50. Typically, 100–200 embryos were plated on 6 cm NGM plates, 4,000–8,000 embryos on 10 cm NGM plates seeded with 10x concentrated bacteria, 15,000 embryos on 15 cm NGM plates seeded with 10x concentrated bacteria.

For collection of starved dauers used for DAF-16 ChIP-seq, wild-type and *daf-16(ar620[daf-16*::*zf1-wrmScarlet-3xFLAG])* animals were grown at 20°C. Worms were developmentally synchronized by sodium hypochlorite treatment. Briefly, eggs were isolated by lysing gravid hermaphrodites in sodium hypochlorite, suspended in M9 buffer, and incubated for 24 h with aeration at room temperature to allow L1 arrest. Starved L1d larvae were plated on NGM plates seeded with HB101. Approximately 15,000 L1d hatchlings were plated on 15 cm NGM plates seeded with 10x concentrated bacteria and maintained at 25°C on Day 1. Animals were re-fed on Day 3, and starved worms were subject to SDS isolation on Day 5. Starved worms were incubated with 1% SDS in water for 30 min with aeration, washed with water, and plated on 15 cm NGM plates to allow viable worms to move out. The surviving worms were collected for ChIP-seq.

### CRISPR/Cas9 mediated genome editing of *C. elegans*

To construct *nhr-23(xk40[nhr-23::aid::3xflag]),* crRNA oHG308 (40 μM, IDT Alt-R CRISPR crRNA), amplified and purified repair template oHG309 (100 ng/μL, initially ordered as an IDT gBlock Gene Fragment), *dpy-10* crRNA (5.6 µM, IDT Alt-R CRISPR crRNA), *dpy-10* repair template (12 ng/µl, IDT Ultramer DNA oligo), tracrRNA (40 µM, IDT Alt-R CRISPR-Cas9 tracrRNA), and Cas9 (15.5 µM, stock at 40 µM in 20 mM HEPES-KOH pH 7.5, 150 mM KCl, 10% glycerol, 1 mM DTT [Dithiothreitol] from Berkeley QB3 MacroLab) were injected into QK159 *nhr-23(xk22[nhr-23::3xflag])* (Patel et al., 2022). All reagents were diluted in IDT duplex buffer. F1 offspring that exhibited Dpy or Rol phenotypes were singled and genotyped for the *nhr-23::AID::3xflag* allele. One line *nhr-23(xk40)* was backcrossed to N2 four times and then to CA1200 *ieSi57II* to generate QK200.

To construct *let-7(scRORE1,2,3)*, crRNA oHG282 and repair template oHG368 (120 ng/µl, IDT Ultramer DNA oligo) were injected as described above into QK198 *let-7(xk39(scRORE1,3))* (Patel et al., 2022). 2 independent lines were backcrossed three times to N2 to generate *let-7(xk45(scRORE1,2,3)* and *let-7(xk58(scRORE1,2,3))*.

To construct *daf-16(xk49 [daf-16::3xflag],* crRNAs oAA1785 and oAA1786 (20 μM each, IDT Alt-R CRISPR crRNA) and repair template oAA1789(100 ng/μL, initially ordered as an IDT gBlock Gene Fragment) were injected as described above into wild type animals. One of the lines homozygous for *daf-16(xk49)* was then injected with crRNA oAA1817 and repair template oAA1818(120 ng/µl, IDT Ultramer DNA oligo) to generate *daf-16::3xflag(N215A R218A H219A*). These animals were further injected with crRNAs oAA1926 and oAA1927 and repair template oAA1928 to generate the full DNA Binding Dead *daf-16(xk48[daf-16::3xflag(DBD/N215A R218A H219A S22A)]* allele.

### Construction of overexpression strains

The oligos used for cloning are listed in Table S3. The plasmid with *col-183p::let-7::let-7 3’ UTR* was generated by cloning the following fragments into house vector pJK585: *col-183* endogenous promoter (1.7 kb region upstream of *col-183* as defined in Shih et al., 2019) and primary *let-7* C05G5.6.2::*let-7* 3’ UTR (891 bp shorter primary *let-7* transcript as defined in Kai et al., 2013 with additional 30 bp sequence downstream of *let-7*). To generate the QK244, the above construct was injected at a concentration of 2 ng/μL along with the co-injection plasmid pJK343 (*myo-2p::mCherry::unc-54 3’ UTR*) at 1.5 ng/μL into VT1777 animals.

The plasmid with *col-183p::nhr-23::3xflag::nhr-23 3’ UTR* was generated by cloning the following fragments into house vector pJK585: *col-183* endogenous promoter (1.7 kb region upstream of *col-183* as defined in Shih et al., 2019) and *nhr-23::3xflag::nhr-23 isoforms b, d, e, f* (3.5 kb region in *nhr-23* gene cloned from QK159 *nhr-23::3xflag; nhr-23(xk22*) (Patel et al., 2022)). To generate the QK245, the above construct was injected at a concentration of 2 ng/μL along with the co-injection plasmid pJK343 (*myo-2p::mCherry::unc-54 3’ UTR*) at 1.5 ng/μL into VT1777 animals.

### Working with AID tagged alleles

Protocols from Zhang et al. (2015) were used to inducibly degrade NHR-23::AID::3xFlag. Briefly, the appropriate volume of auxin (3-indole acetic acid; Sigma-Aldrich I2886) dissolved in 100% ethanol was added to autoclaved liquid NGM. Plates were seeded with HB101, maintained in the dark, and used within one month. Worms were transferred to auxin plates at the appropriate developmental time points. For experiments using *nhr-23::GFP::flag::AID* (Ragle et al., 2020), dauers were incubated in 1 mM auxin in M9 buffer with aeration for 2 h.

### Quantitative fluorescence microscopy

Animals were immobilized with 50 mg/mL levamisole and mounted on 2% agarose pads. Worms were imaged using a Zeiss AxioImager M2 or D2 microscope with an ORCA-Flash 4.0 LT or Axiocam 807 camera at 40x or 63x magnification and processed using Zeiss Zen2 or Zen 10 software. Representative images correspond to the best-focused optical section. Fluorescence intensity was quantified using FIJI. For each animal, fluorescence from three or more hypodermal cells was measured and averaged.

### RNA extraction

Hypochlorite-prepped embryos were directly plated on HB101. Animals were assessed for dauer morphology and 500–1,000 dauers were collected at 52 h at 24°C in TRI Reagent (ThermoFisher Scientific AM9738). For L2d samples, animals were collected beginning at 30 h at 24°C. Following five freeze-thaw cycles, 1-bromo-3-chloropropane was added and RNA in the aqueous phase was precipitated with isopropanol for 2 h at −30°C. Samples were centrifuged at 21,000 x g for 30 min at 4°C to pellet RNA. Pellets were washed three times with 75% ethanol and resuspended in nuclease-free water.

### RT-qPCR

cDNA synthesis for primary and precursor *let-7* transcripts was performed using SuperScript III Reverse Transcriptase (Invitrogen 18080044). A total of 250 ng RNA was used for cDNA synthesis using an Eppendorf Mastercycler Pro S6325. Quantitative PCR for *pri-let-7, nhr-23, mlt-10* and *eft-2* was performed using Absolute Blue SYBR Green (Thermo Scientific AB4166B) on a CFX63 Real Time System Thermocycler (Biorad) using custom primers. Cycle numbers for *pri-let-7, nhr-23* and *molt-10* were normalized to respective cycle numbers for *eft-2*. Two biological replicates with two technical replicates were performed unless otherwise stated. Statistical significance was evaluated using a two-tailed Student’s t-test.

For mature microRNAs, TaqMan assays were performed using probes from Applied Biosystems. A total of 100 ng RNA was used for reverse transcription with Multiscribe Reverse Transcriptase (ThermoFisher Scientific 4311235). Quantitative PCR for *let-7, mir-48, mir-84, mir-241*, and U18 were performed using TaqMan Universal Master Mix, No AmpErase UNG (ThermoFisher Scientific 4324018) on a CFX63 Real Time System Thermocycler (Biorad). TaqMan probes and Assay IDs were: *let-7* (000377), *mir-48* (000208)*, mir-84* (463262)*, mir-241* (000249), and U18 (001764) (ThermoFisher Scientific). The cycle numbers for *let-7, mir-48, mir-84, mir-241* were normalized to respective cycle numbers for U18. Three biological replicates with two technical replicates were performed unless otherwise stated. Statistical significance was evaluated using a two-tailed Student’s t-test.

### RNA-seq

RNA samples were checked for quality/quantity on Fragment Analyzer and Qubit prior to library prep. NEBNext rRNA Depletion Kit v2 (NEB # E7400) and NEBNext Ultra II RNA Library Prep Kit for Illumina (Cat# E7765) were used to generate libraries. The Fragment Analyzer was used for quality control of the libraries to ensure adequate concentration and appropriate fragment size. The resulting library insert size was 200bp-500bp with a median size around 300bp. Libraries were barcoded using unique dual indexing (E6440S) and pooled for sequencing. Pooled libraries were sequenced on an Illumina® NovaSeq6000 instrument using standard protocols for SP 2x150bp Paired end sequencing. FASTQ files were generated using bcl-convert v3.7.5.

### Analysis of RNA-seq

Read quality was assessed using FastQC (http://www.bioinformatics.babraham.ac.uk/projects/fastqc). Adapter sequences and low-quality bases were trimmed using Trim Galore! v0.6.6 (http://www.bioinformatics.babraham.ac.uk/projects/trim_galore/) with parameters – length 36. Trimmed reads were aligned to the *C. elegans* WBCel235 reference genome using STAR (Dobin et al., 2013) with default parameters. Final BAM files were sorted and indexed using samtools for downstream analyses (Li et al., 2009). For every transcript in each replicate, FPKM values across all conditions were normalized to the average FPKM value in the *nhr-23(xk40); ieSi57; daf-7(e1372)* 0 mM auxin condition to generate heatmaps representing relative expression values. No reads mapping to the primary transcripts of *mir-48, mir-84, mir-241* were detected and hence are omitted from the heatmaps in Figure 4A and Figure S9B. The Python library Seaborn (https://seaborn.pydata.org/) were used to generate cluster maps to define Categories 1, 2, and 3 with the expression profiles of the CCG genes. *nhr-23* targets that binned into the same clusters as the CCGs were then added to Categories 1, 2, and 3. The relative values across all the replicates and all the genes were averaged to generate the metaplots in Figure 4A.

### Western blot

Samples were collected in 2x Novex SDS loading buffer and lysed by incubating three times at 95°C for 5 min followed by 4°C for 5 min. Samples were resolved on 8% or 10% Novex Tris-glycine SDS gels and transferred to PVDF membranes (Millipore). Primary antibodies used were mouse anti-Flag (Sigma–Aldrich F1804, 1:1000 dilution), rabbit anti-gamma tubulin (Sigma–Aldrich T1450, 1:2000) and rabbit anti-H3 (Abcam Ab12079, 1:5000). Secondary antibodies used were sheep anti-mouse (GE Healthcare NA931, 1:1000), and goat anti-rabbit (Jackson Laboratories 111035045, 1:2000). Signal detection was performed using Amersham ECL Prime (GE Healthcare, RPN2232) or Pierce ECL (ThermoFisher, 32209) and visualized using a Bio-Rad ChemiDoc Touch system.

### Seam cell counting

Hypochlorite-prepared embryos were hatched for 24 h in M9 buffer with nutation. L1-arrested animals were plated on HB101 at 25°C. Animals were scored 40– 44 h after plating. Worms were immobilized in 50 mg/mL levamisole on 2% agarose pads. Seam cells expressing *Pscm::GFP* were counted using a Zeiss Axio Zoom V16 fluorescence stereomicroscope.

### Chromatin immunoprecipitation for ChIP-seq and ChIP-qPCR

Animals starved at 25°C were collected as ∼500μL packed pellets in M9. Worms were crosslinked in 12mL of 2.6% (v/v) formaldehyde for 30 min at room temperature. Crosslinking was quenched by addition of 600 µL 2.5 M glycine for 5 min. Samples were then washed three times in water and flash-frozen in liquid nitrogen. Frozen pellets were ground using a Retsch MM400 CryoMill for two 1-min cycles at 30 Hz in liquid nitrogen-cooled stainless steel cryomill chambers. The resulting powder was resuspended and further lysed in 2 mL of RIPA buffer (1x PBS, 1% (v/v) NP40, 0.5% sodium deoxycholate, and 0.1% SDS), supplemented with HALT protease and phosphatase inhibitor cocktail (ThermoFisher Scientific) for 10 min at 4°C.

To shear the chromatin, samples were sonicated in a Bioruptor Pico (Diagenode) for three cycles of 3 min each (30 s ON/30 s OFF cycles) at 4 °C. A 20 µL aliquot of the sample was treated with Proteinase K for 10 min and then subjected to phenol chloroform extraction, as described below. The concentration of the aliquot was determined using a Qubit Fluorometer 3.0 (Invitrogen). Based on the initial concentration of the aliquot, the chromatin sample was diluted to 20–30 ng/µL. To check the extent of shearing, the same aliquot was run on an agarose gel. The sample was processed further only if the DNA smear was centered around 200 bp.

Of the total amount of chromatin that remained, 10% was used as the input sample (stored at 4°C) and 90% was subjected to immunoprecipitation. Every 10 μg of chromatin was incubated with 2 µg of mouse M2 anti-Flag monoclonal antibodies (Sigma-Aldrich) overnight at 4°C on a nutator. Samples were then incubated with 1.5 mg of affinity-purified sheep anti-mouse IgG antibodies covalently attached to superparamagnetic Dynabeads M-280 (Invitrogen) for 2 h at 4°C. Thereafter, complexes bound to the beads were separated three times from the supernatant and washed in 800 µL LiCl buffer (100 mM Tris-HCL pH 7.5, 500 mM LiCl, 1% (v/v) NP40, and 1% sodium deoxycholate). The resulting immunoprecipitates were de-crosslinked by incubation with 80 μg of proteinase K in 400 μL of worm lysis buffer (100 mM Tris-HCL pH 7.5, 100 mM NaCl, 50 mM EDTA, and 1% SDS) at 65 °C for 4 h; input samples underwent the same treatment in parallel. Residual proteins were removed from both IP and input samples by phenol-chloroform extraction. Briefly, 400 µL of phenol-chloroform-isoamyl alcohol pH 8.0 (Sigma-Aldrich) was added to each sample. Samples were vortexed vigorously and centrifuged at 15,000 x g for 5 min at 4 °C. The aqueous phase top layer was transferred to a new tube and DNA was precipitated by incubation with 1 mL of 0.3 M ammonium acetate (Sigma-Aldrich) in ethanol for 1 h at −30°C. The resulting DNA pellet was washed twice with 100% ethanol and resuspended in Tris-EDTA, pH 8.0. Prior to use as a template for qPCR, the entire DNA sample was treated with RNase A for 1 h at 37°C. ChIP-seq libraries were prepared using the Ovation Ultralow System V2 kit (NuGEN, 0344NB-A01) according to the manufacturer’s protocol, with 15 PCR amplification cycles. Libraries were sequenced on an Illumina NovaSeq 6000 instrument using paired-end 50 bp reads.

Quantitative PCR for promoter regions of interest was performed using Absolute Blue SYBR Green (Thermo Scientific) on a CFX96 Real Time System Thermocycler (BioRad) according to the manufacturer’s instructions. The Ct value for each IP sample was first normalized to the Ct value for the corresponding input sample. The log_2_-transformed fold-change values for samples derived from *daf-16::3xflag* were then normalized to the respective N2 sample. Two technical replicates were performed for each amplicon of interest, as specified in the corresponding figure legends, in two biological replicates.

### Analysis of ChIP-seq data

All samples were sequenced across two lanes, and resulting FASTQ files were merged prior to downstream processing. Read quality was assessed using FastQC (http://www.bioinformatics.babraham.ac.uk/projects/fastqc). Adapter sequences and low-quality bases were trimmed using Trim Galore! v0.5.0 (http://www.bioinformatics.babraham.ac.uk/projects/trim_galore/) with parameters -- quality 25 and –stringency 3. Trimmed reads were aligned to the *C. elegans* WBCel235 reference genome using Bowtie2 (Langmead and Salzberg 2012) with default parameters. Alignments were filtered using samtools (Li et al., 2009) to remove unmapped, secondary, and low-quality alignments. PCR duplicates were identified with Picard MarkDuplicates 2.22.1 (http://broadinstitute.github.io/picard/) and subsequently removed. Final BAM files were sorted and indexed using samtools for downstream analyses.

Binding profiles for DAF-16 were identified using MACS2 at a p value threshold of 0.05 (Zhang et al., 2008). Fold-enrichment tracks were generated with MACS2, with blacklisted regions removed, and converted into bigWig format using the bedGraphToBigWig command-line utility from the UCSC Genome Browser kentUtils package (Kent et al., 2002). Coverage tracks were visualized in the Integrative Genomics Viewer (Robinson et al., 2011). Peaks from biological replicates for each genotype were intersected using BEDTools (Quinlan and Hall, 2010), required at least 10% reciprocal overlap, and overlapping regions were retained as consensus summit regions. From these consensus regions, those showing at least 2-fold higher signal in starved dauers expressing *daf-16::mScarlet::3xflag* compared to wild-type starved dauers were retained for further analysis. Retained summit regions were annotated to the nearest gene using HOMER (Heinz et al., 2010).

### Motif analysis

DNA sequence motifs near DAF-16 binding sites were identified using the MEME suite (Bailey et al., 2009). Using BEDTools getfasta (Quinlan and Hall, 2010), FASTA files containing 200 bp genomic sequences centered on DAF-16 binding sites were extracted from the *C. elegans* WBcel235 reference genome. Motif discovery was performed independently on positively and negatively regulated target gene set using MEME, with the number of motifs to identify set to 10 and motif widths constrained to 6-10bp. Motif enrichment and positional distribution relative to binding sites were further assessed using MEME-ChIP.

### Gene Ontology analysis

Genes that were differentially expressed (>2-fold change in *daf-16(-)* versus control dauers and FDR<0.05) in Wirick et al., 2021 and were predicted to be DAF-16 targets were used for Gene Ontology (GO) analysis (Ashburner et al., 2000; The Gene Ontology Consortium 2005). Positively and negatively regulated targets were processed separately using PANTHER (Mi et al., 2019).. The PANTHER Overrepresentation Test was performed using the GO Ontology database (DOI: 10.5281/zenodo.6799722 Released 2022-07-01). The GO biological Process Complete dataset was used as the annotation data set. Fisher’s exact test was used to calculate FDR.

### Quantification and statistical analyses

Python 3, GraphPad Prism v6.0h and Microsoft Excel were used for all statistical analyses. Sample sizes for all experiments, statistical tests, and resulting p values are provided in the corresponding figure legends.

## Data Availability

The data discussed in this publication have been deposited in NCBI’s Gene Expression Omnibus (Edgar *et al*., 2002) and are accessible through GEO Series accession number GSE338292 (ChIP-seq) and GSE338973 (RNA-seq).

## Acknowledgement

Library prep and Illumina sequencing for RNA-seq were conducted at the Integrated Genomics Center, Johns Hopkins Department of Genetic Medicine, Baltimore, MD. We thank Dr. Luisa Cochella, Dr. Oliver Hobert, Dr. Jordan Ward and Dr. Iva Greenwald for sharing strains and reagents. We thank Amy Liang and Shreya Singireddy for their work in construction of strains and optimizing reagents. We acknowledge members of the Kim and Karp labs for insightful discussions and helpful suggestions. This work was supported by the National Institute of Health (NIH) grant R01 HD109667-01 to J.K.K., NIH grant R15 GM150082-01 to XK. S.E.J. is an Investigator of the Howard Hughes Medical Institute.

---

**Fig. S1.**
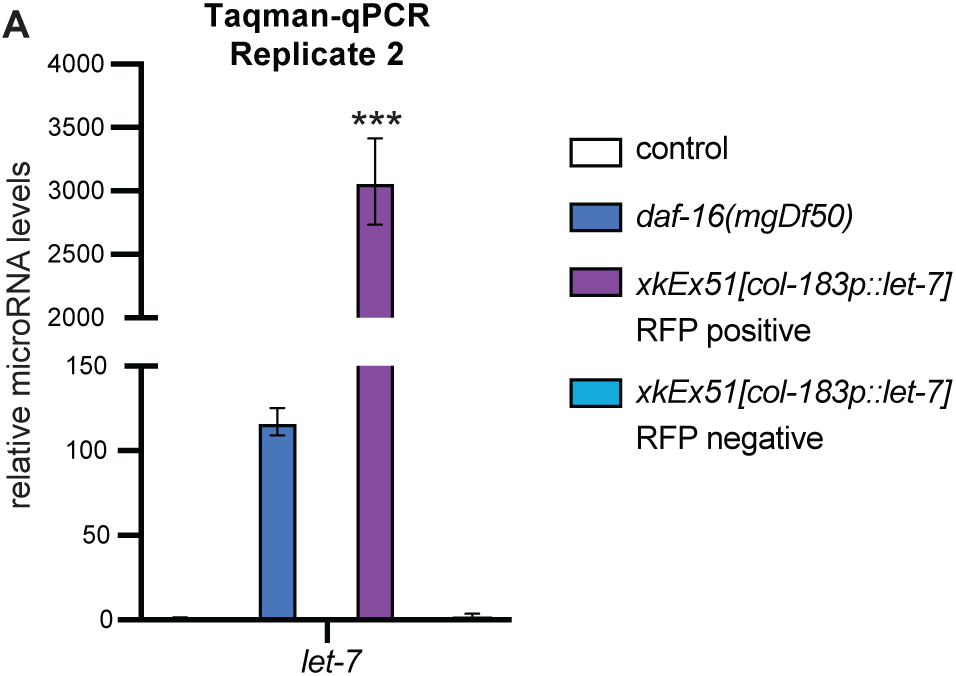
Validation of overexpression of *let-7* during dauer. **A.** Taqman qPCR measuring *let-7* levels during dauer in the indicated genotypes with *daf-7(e1372); maIs105[col-19p::gfp]* in the background. Expression was normalized to U18 and then to the value in control *dauers*. Bars represent mean of two technical replicates and error bars represent standard deviation. P values were calculated using Student’s t-test. *** p<0.001.

**Fig. S2.**
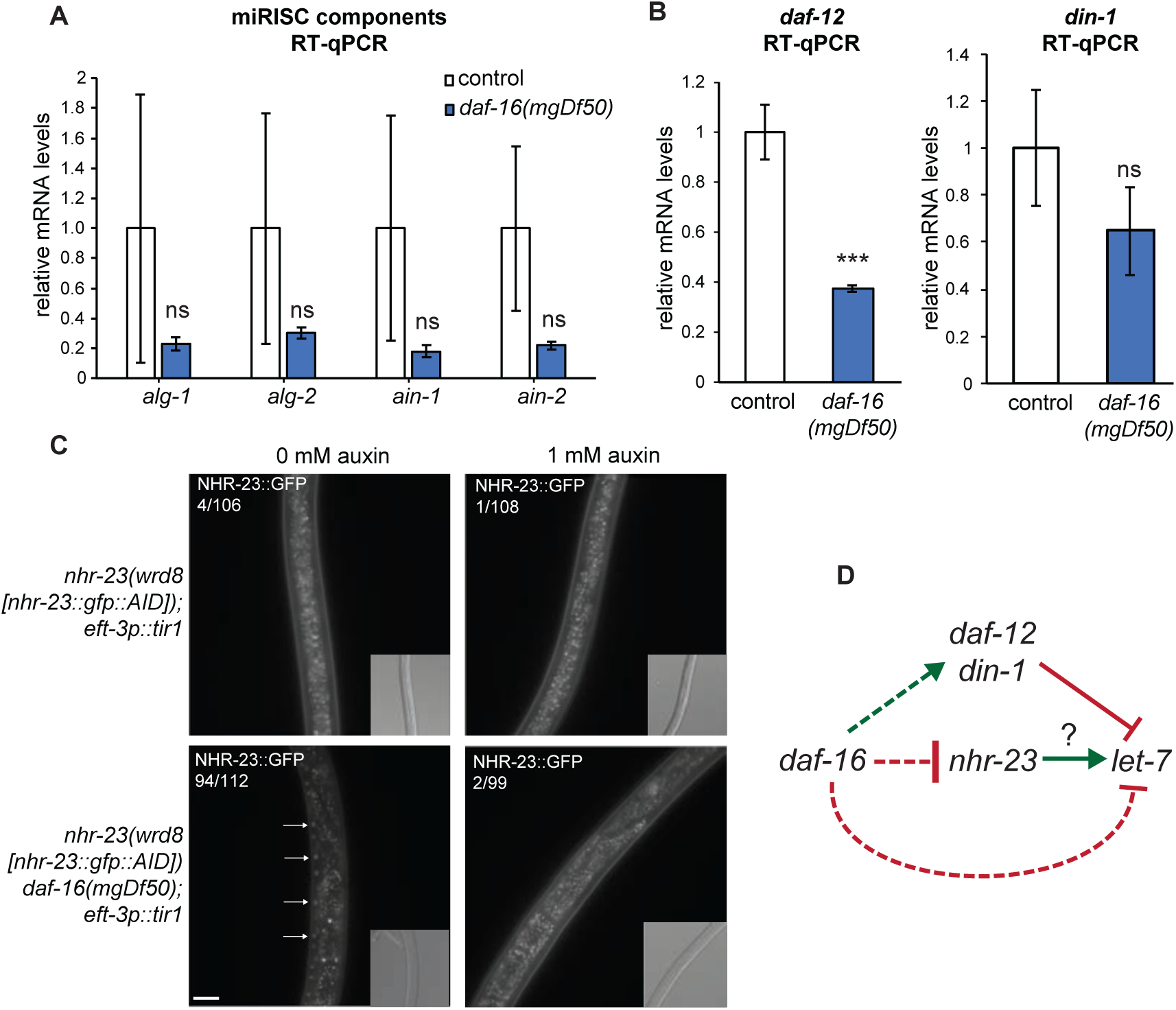
*nhr-23* is upregulated and *daf-12* is downregulated in *daf-16(mgDf50); daf-7(e1372)* dauers. All strains in this figure have *daf-7(e1372)* in the background. **A, B.** RT-qPCR measuring expression of **A**. indicated components of the miRISC and **B.** *daf-12* and *din-1* in control and *daf-16(mgDf50)* dauers. Bars represent the mean of three biological replicates and error bars represent standard deviation. P values were calculated using Student’s t-test. n.s., non significant.***p<0.001. **C.** Representative images showing NHR-23::GFP expression in the indicated genotypes and auxin conditions. Strains have *ieSi57[eft-3p::tir1].* The number of animals expressing NHR-23::GFP::AID is indicated. Arrows indicate seam cell nuclei. Images of 0 mM auxin condition on the left are reproduced from Figure 2E. Scale bar = 20μm **D.** Schematic summarizing the genetic interactions established in this figure and in Figure 2A–E. Dashed lines indicate possible involvement of additional factors. The question mark indicates the pathway investigated in Figure 2F–H.

**Fig. S3.**
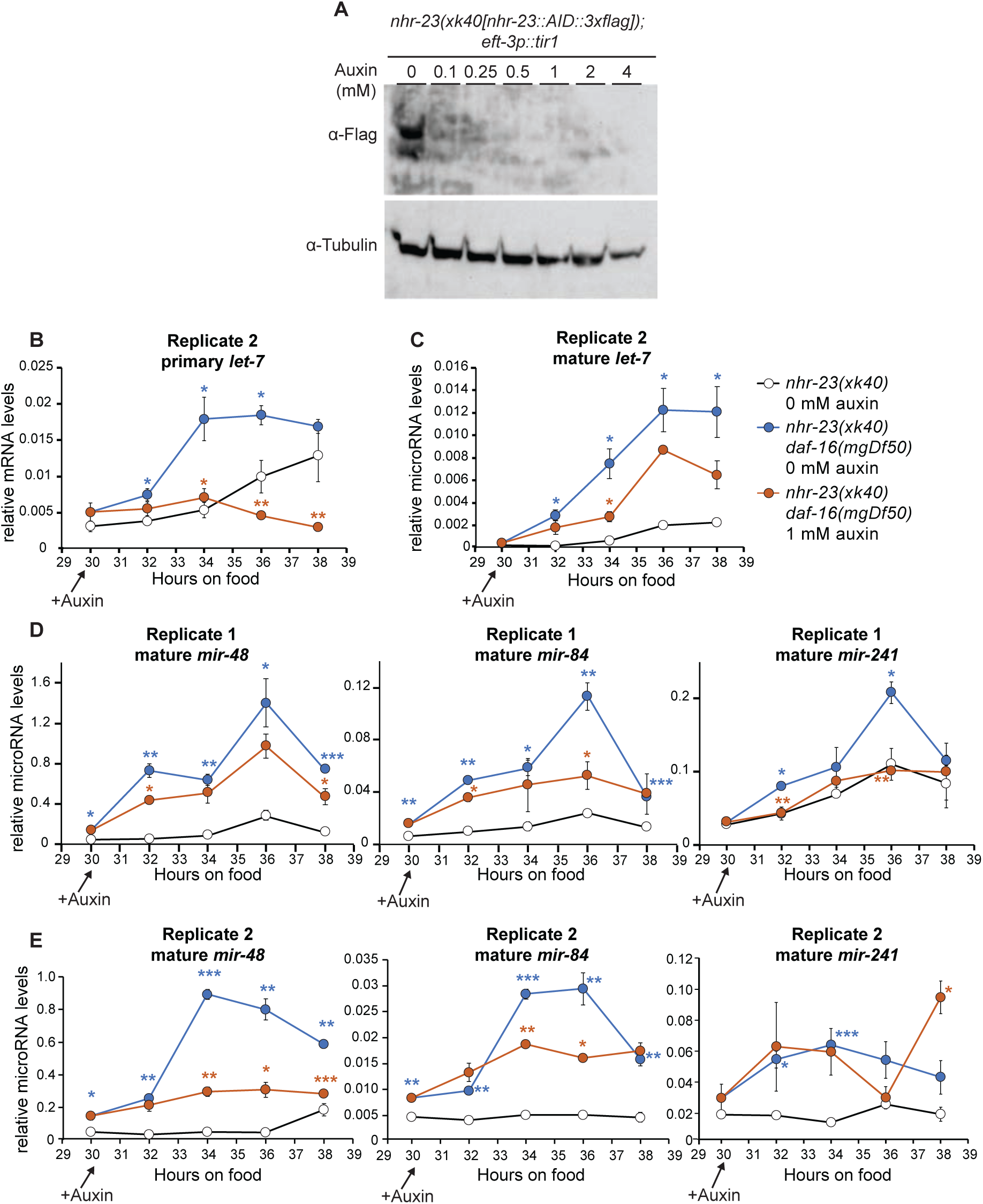
NHR-23 mediates upregulation of the *let-7* family of microRNAs in *daf-16(mgDf50); daf-7(e1372)* dauers. **A.** Western blot showing NHR-23::AID::3xFlag levels in starved dauers after exposure to the indicated concentrations of auxin for 2 h. **B–E.** All strains have *daf-7(e1372); ieSi57[eft-3p::tir1]* in the background. RT-qPCR measuring primary *let-7* levels **(B)** and Taqman qPCR measuring mature *let-7* **(C)***, mir-48, mir-84* and *mir-241* levels **(D-E)** in the indicated genotypes. Worms were exposed to 0 mM or 1 mM auxin at 30 h. Each data point represents the mean of two technical replicates and error bars represent standard deviation. Blue asterisks indicate comparisons between *nhr-23::AID,* 0 mM and *nhr-23::AID daf-16(mgDf50),* 0 mM auxin. Orange asterisks indicate comparisons between *nhr-23::AID daf-16(mgDf50)* 0 mM auxin and *nhr-23::AID daf-16(mgDf50),* 1 mM auxin. P values were calculated using Student’s t-test. *p<0.05, **p<0.01, ***p<0.001.

**Fig. S4.**
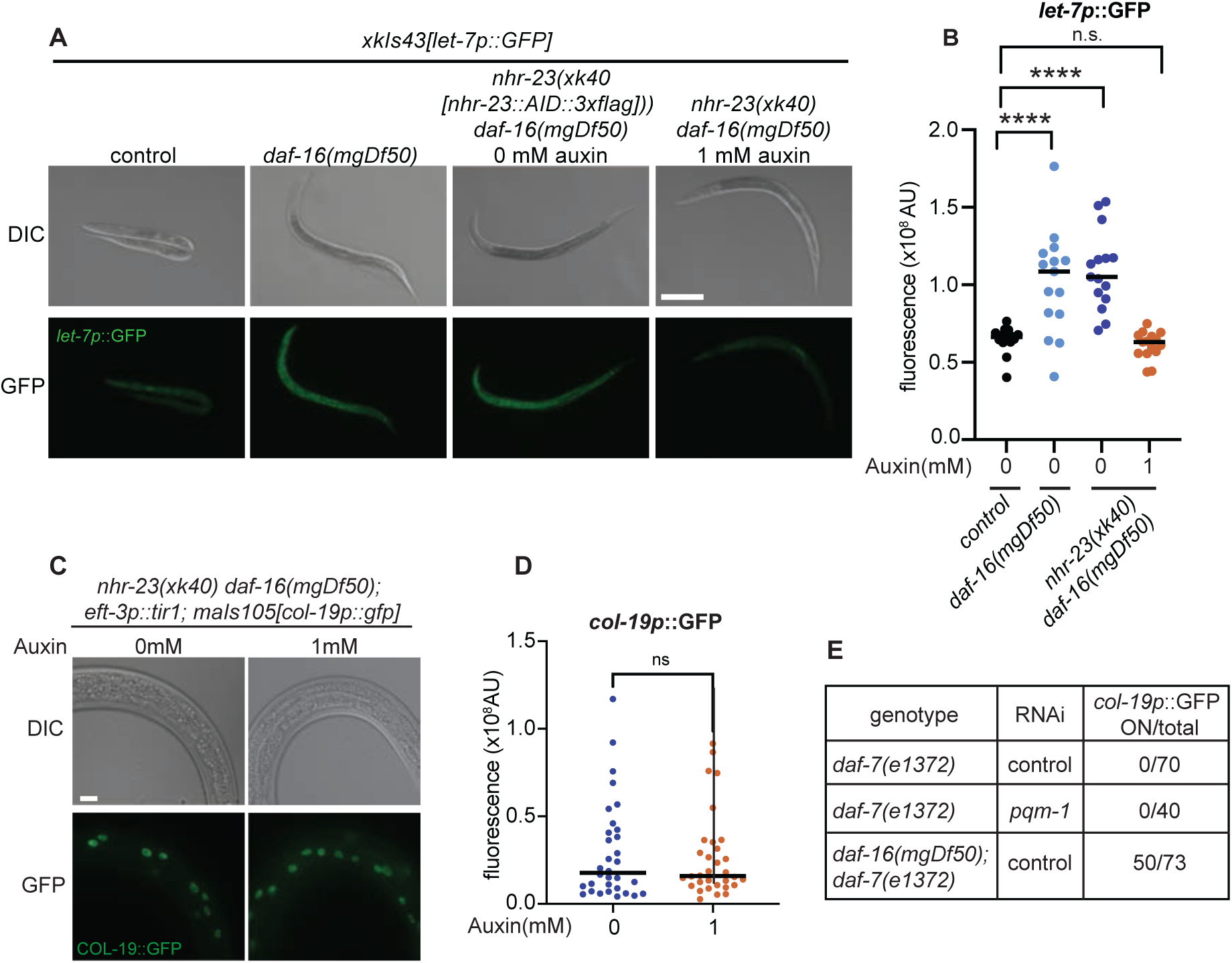
NHR-23 mediates upregulation of *let-7* transcription in *daf-16(mgDf50); daf-7(e1372)* during L2d. All strains have *daf-7(e1372).* **A.** Representative images of indicated genotypes at 38 h at 24°C. Animals were transferred to plates containing 0 mM or 1 mM auxin at 28 h. Scale bar = 100μm. **B.** Quantification of images shown in A. Each data point represents the mean fluorescence intensity from at least three hypodermal cells per animal. N = 14–20. P values were calculated using the Mann Whitney test. n.s., not significant; ****p<0.0001. **C.** Representative images of *col-19p*::GFP expression in dauers of indicated genotypes exposed to 0 mM or 1 mM auxin for 2 h. Scale bar = 10μm **D.** Quantification of images shown in C. Each data point represents the mean fluorescence intensity from at least three hypodermal cells per animal. N = 14–20. P values were calculated by using the Mann Whitney test. n.s., not significant; ****p<0.0001. **E.** Table showing number of dauers expressing *col-19p*::GFP in indicated conditions.

**Fig. S5.**
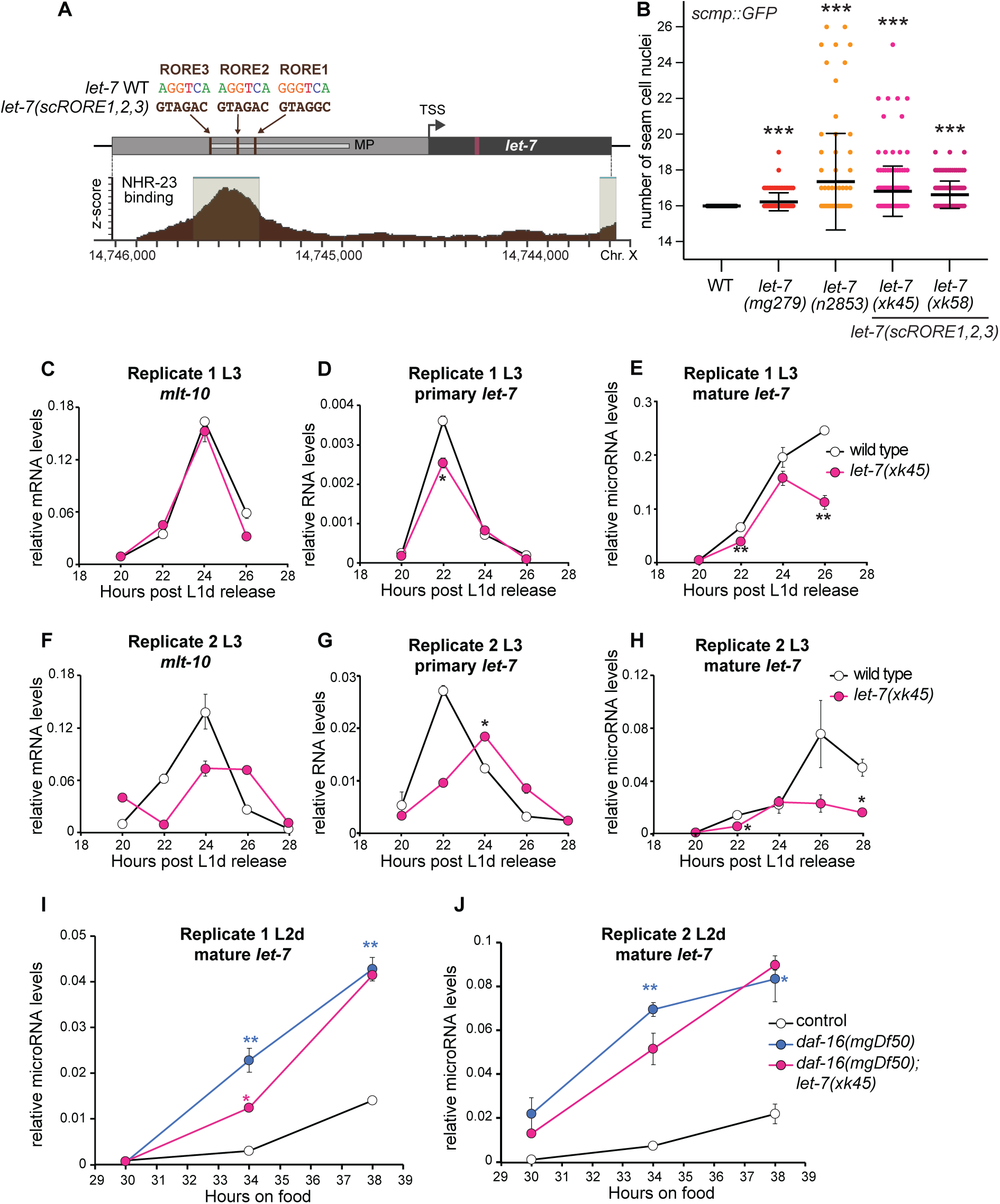
ROREs mediate NHR-23 binding to the *let-7* promoter to promote transcription during continuous development and L2d. **A.** Schematic of the *let-7* promoter showing ROR binding element (RORE) sequences in wild-type and *let-7(scRORE1,2,3)* alleles. MP indicates the minimal promoter from Kai et al., 2013. TSS indicates the transcription start site. Adapted from Patel et al., 2022. **B.** Number of seam cell nuclei in adults of the indicated genotypes. Each data point represents a single animal. Thick line indicates the mean of the distribution and error bars represent standard deviation. N = 67-169 for each genotype. P values were calculated using Student’s t-test. ***p<0.001. **C–H.** Expression analysis during the L3 stage. RT-qPCR measuring *mlt-10* **(C,F)** and primary *let-7* **(D,G)** levels. **E, H.** Taqman qPCR measuring mature *let-7* expression during L3. **I, J**. Strains have *daf-7(e1372).* Taqman qPCR measuring mature *let-7* levels during L2d. Data points represent the mean of two technical replicates and error bars represent standard deviation. P values were calculated using Student’s t-test. *p<0.05, **p<0.01. In G–H, blue asterisks indicate comparisons between control and *daf-16(mgDf50)*, while pink asterisks indicate comparisons between *daf-16(mgDf50)* and *daf-16(mgDf50); let-7(xk45(scRORE1,2,3))* animals. In D and G, comparisons were made between the amplitudes of the two gene expression curves.

**Fig. S6.**
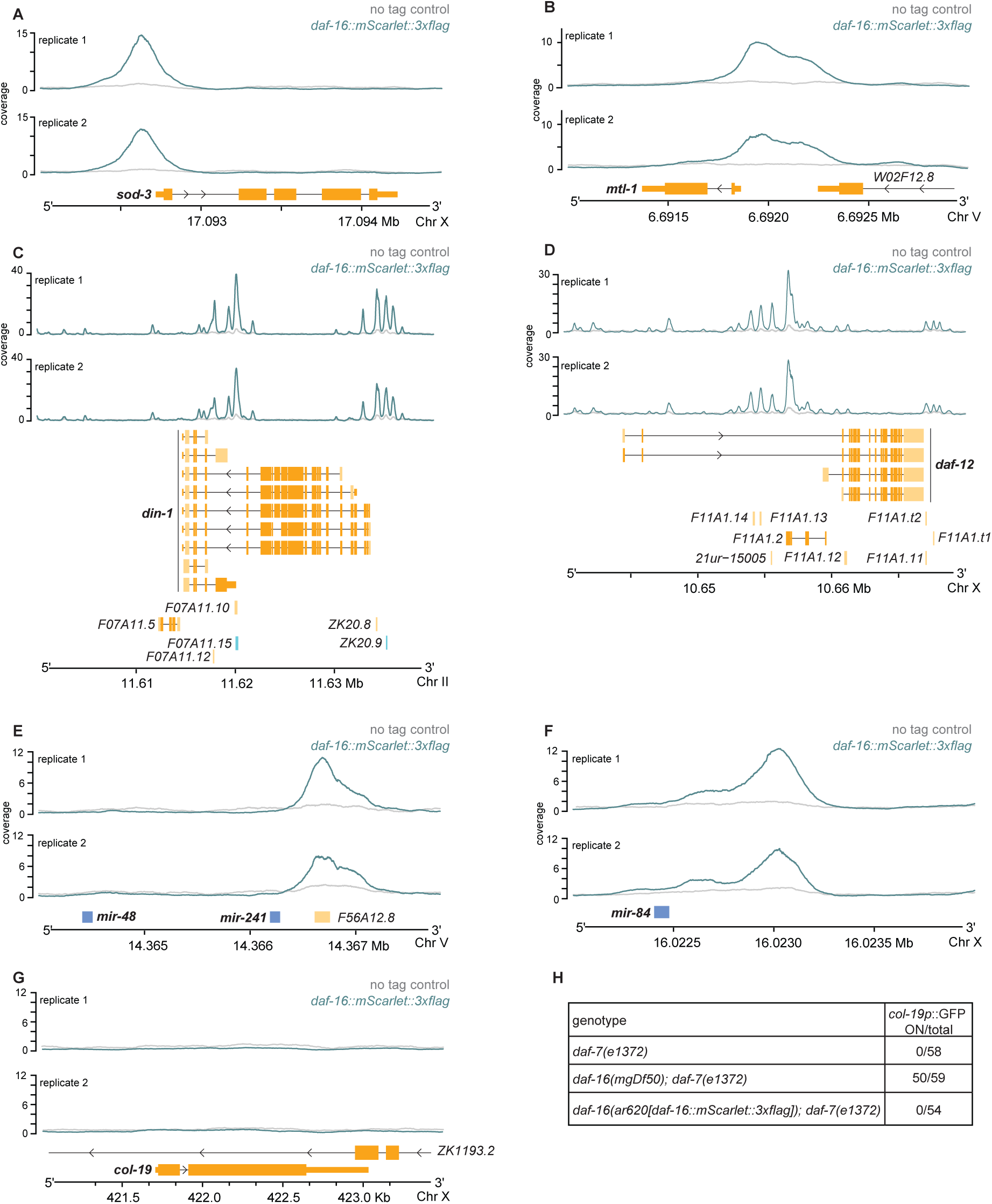
DAF-16 binds upstream of several genes during dauer. ChIP-seq tracks for N2 and *daf-16::mScarlet::3xflag* dauers showing DAF-16 binding summits upstream of **A.** *sod-3,* **B.** *mtl-1*, **C.** *daf-12,* **D.***din-1,* **E.** *mir-241*, and **F.** *mir-84*. **G.** DAF-16 binding summits were not detected upstream of *col-19*. **H.** Strains contain *maIs105[col-19p::gfp]*. Number of dauers expressing *col-19p*::GFP is indicated.

**Fig. S7.**
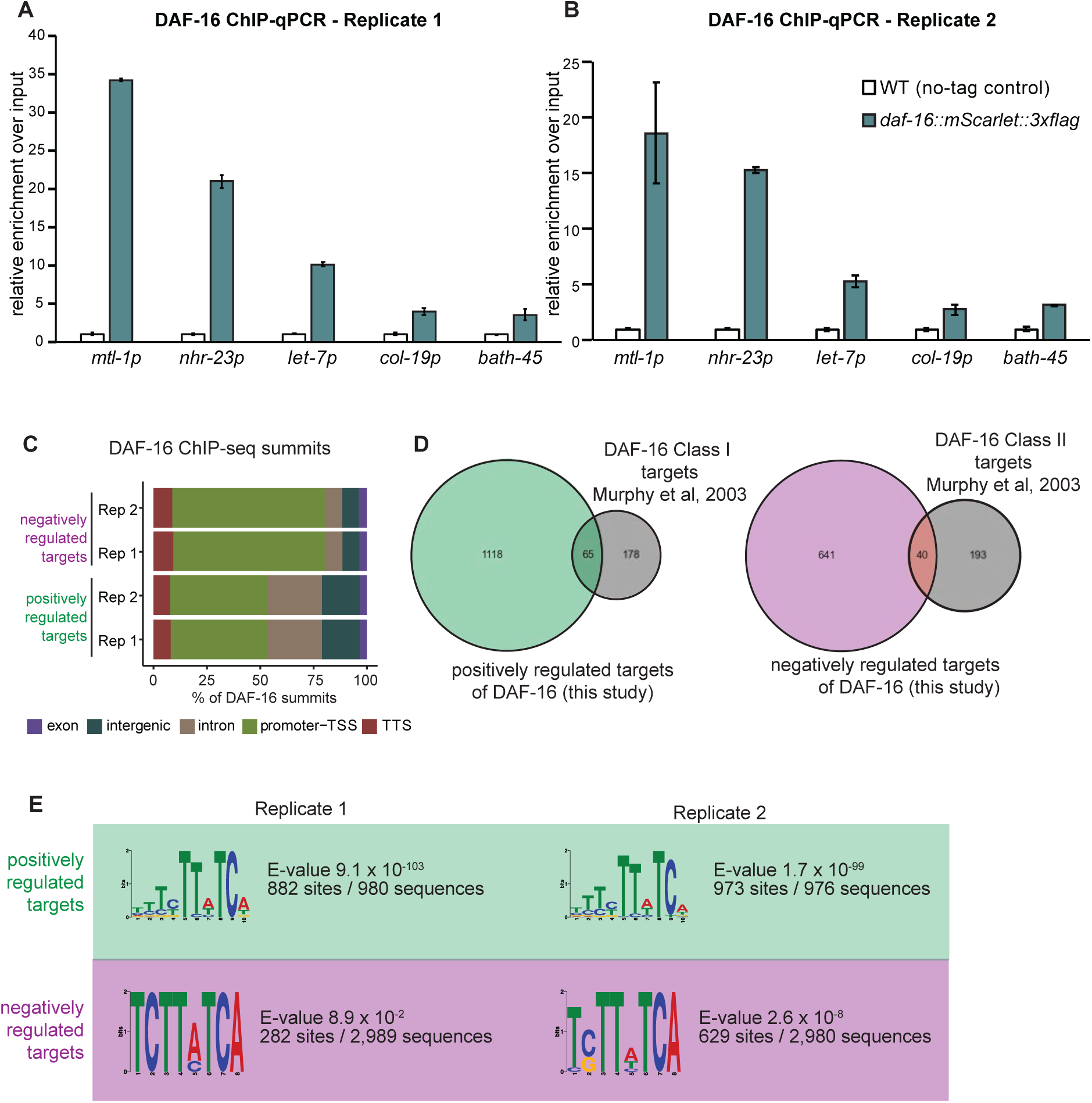
Analysis of DAF-16 ChIP-seq data. **A, B.** ChIP-qPCR validation of DAF-16 enrichment at promoter region of *nhr-23 and let-7* during dauer. Two biological replicates are shown. Bars represent the mean of two technical replicates from one biological replicate and error bars represent standard deviation. The known DAF-16 target *mtl-1* and a heterochromatinized gene *bath-45* were used as positive and negative controls, respectively. **C.** Distribution of DAF-16 consensus binding summits across annotated genomic regions. **D.** Overlap between DAF-16 targets identified in this study and targets previously reported by Murphy et al., 2003. **E.** MEME-ChIP analysis of positively and negatively regulated DAF-16 targets.

**Fig. S8.**
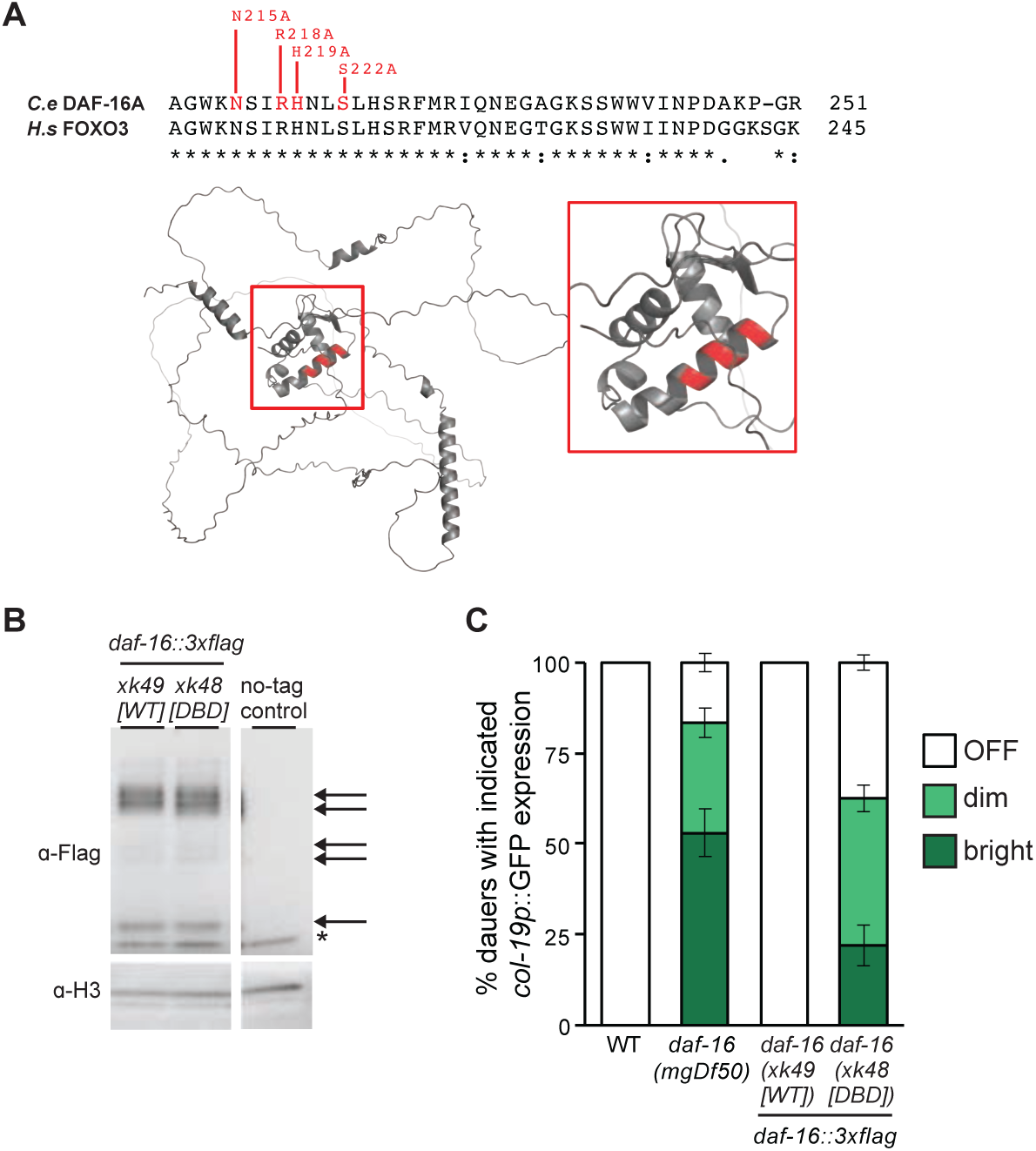
*daf-16::3xflag(DBD)* is a reduced function allele of *daf-16*. **A.** Top: Alignment of the FOXO domain of *C. elegans* DAF-16A and human FOXO3. Labeled residues are required for DNA binding of human FOXO3 (Tsai et al., 2007). Bottom: Predicted structure of *C. elegans* DAF-16A generated using AlphaFold. Conserved residues mutated to alanine in the DNA-binding defective allele *daf-16*(*xk48[daf-16::3xflag(DBD)])* are highlighted in red. **B.** Western blot of DAF-16::3xFLAG in *daf-7(e1372); daf-16*(*xk49[daf-16::3xflag(WT)])* and *daf-7(e1372); daf-16*(*xk48[daf-16::3xflag(DBD)])* dauers. Arrows indicate bands corresponding to FLAG-tagged DAF-16 isoforms. Asterisk indicates a non-specific band. **C.** Percentage of *daf-7(e1372)* dauers exhibiting *col-19p*::GFP expression in the indicated genotypes at 52 h at 25°C. N = 200–300 animals per genotype.

**Fig. S9.**
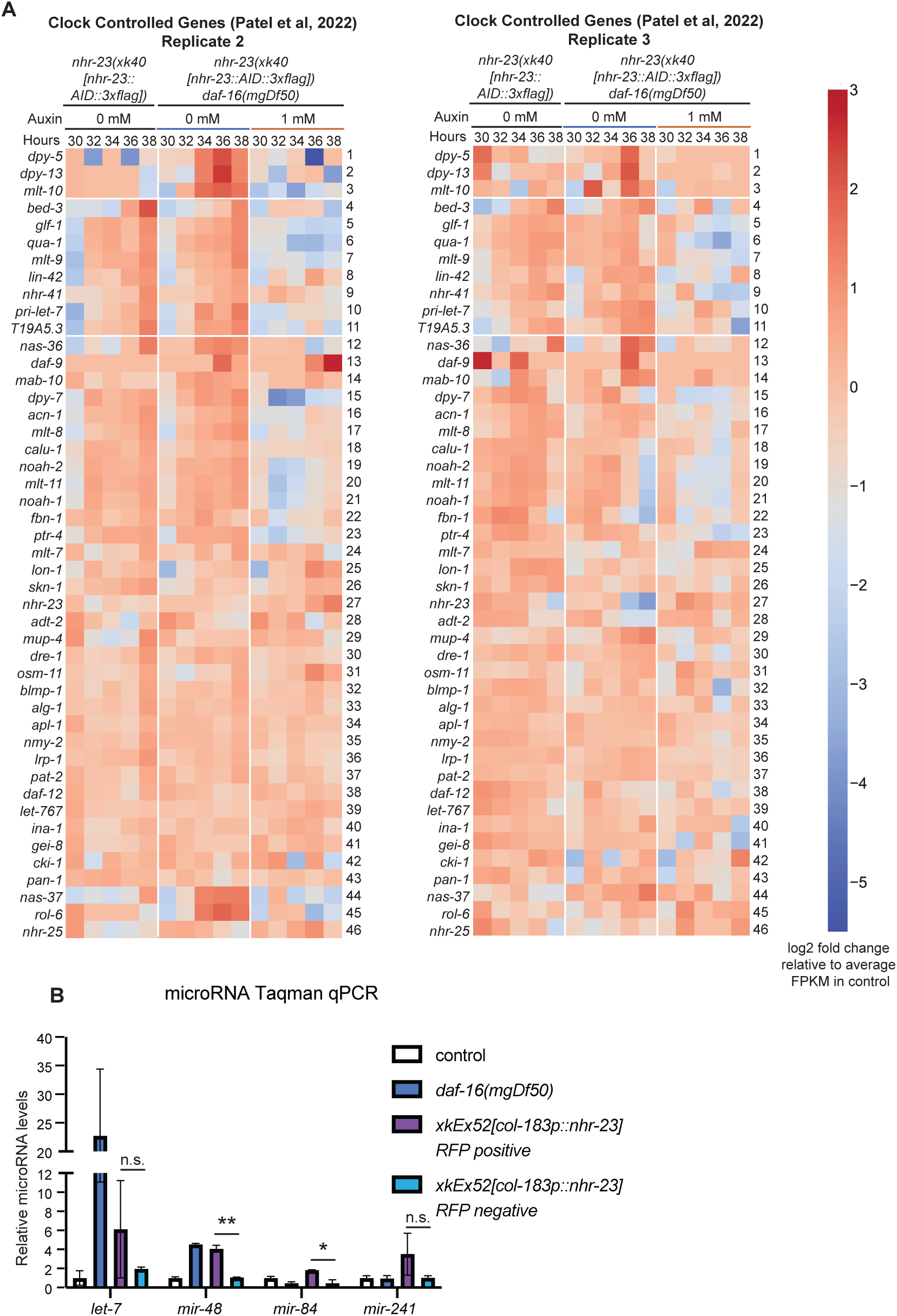
Suppression of *nhr-23* by DAF-16 affects expression of genes associated with continuous development. **A.** Heat maps showing expression profiles of Clock Controlled Genes (CCGs) in the indicated genotypes during development L2d development. Strains have *daf-7(e1372); ieSi57*. Animals were transferred to plates containing 0 mM or 1 mM auxin at 30 h and maintained at 24°C. **B.** Taqman qPCR measuring mature *let-7, mir-48, mir-84* and *mir-241* levels in the indicated genotypes during dauer. Strains have *daf-7(e1372); maIs105[col-19p::gfp]*. P values were calculated using Student’s t test. *p<0.05, **p<0.01; n.s., not significant.

**Fig. S10.**
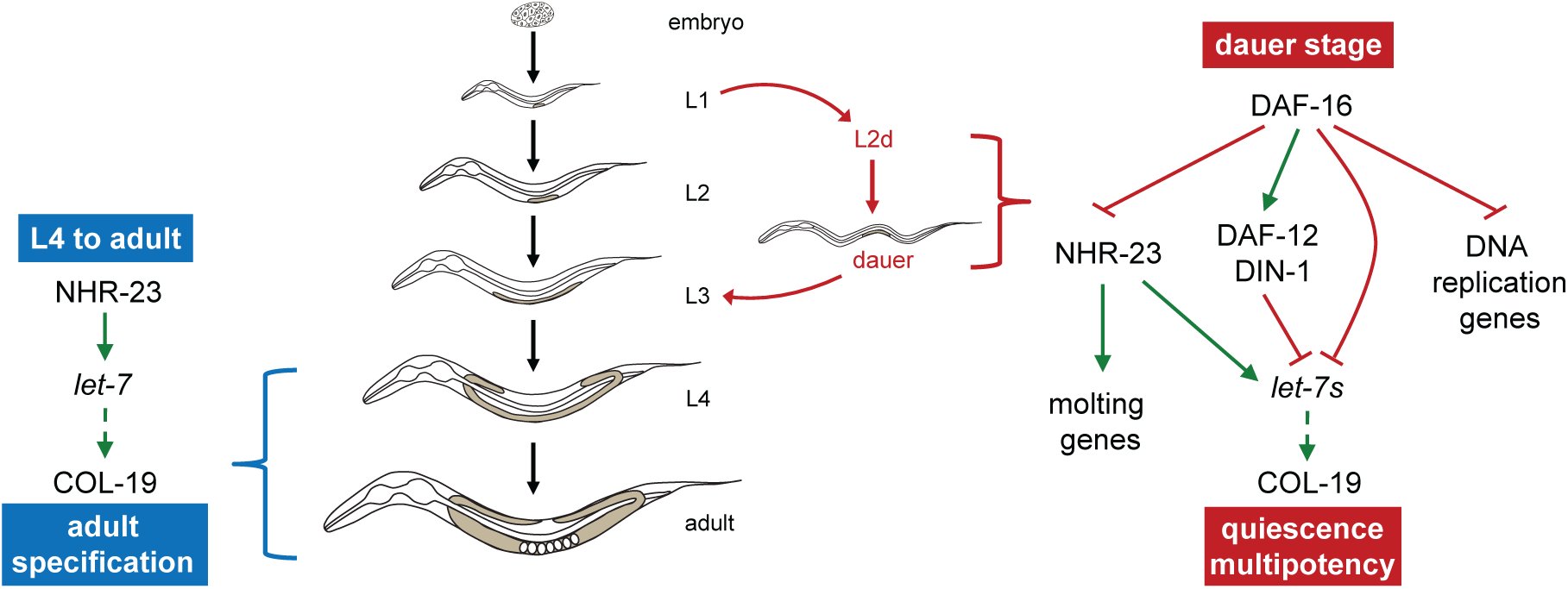
Model: DAF-16 modulates NHR-23 and other pro-growth genes to establish quiescence during dauer. DAF-16 binds upstream of *nhr-23* and suppresses its expression. Because NHR-23 is an essential transcription factor for molting genes (Kouns et al., 2011) and for *let-7* transcription (Patel, Galagali et al., 2022), DAF-16-mediated repression of *nhr-23* may contribute to induction of developmental quiescence. DAF-16 may also promote expression of *daf-12* and *din-1*, which together inhibit *let-7* transcription during dauer (Hammell et al., 2009). In addition, DAF-16 may directly repress transcription of *let-7* and other pro-growth genes, thereby establishing and reinforcing quiescence during dauer.

**Table S1.**
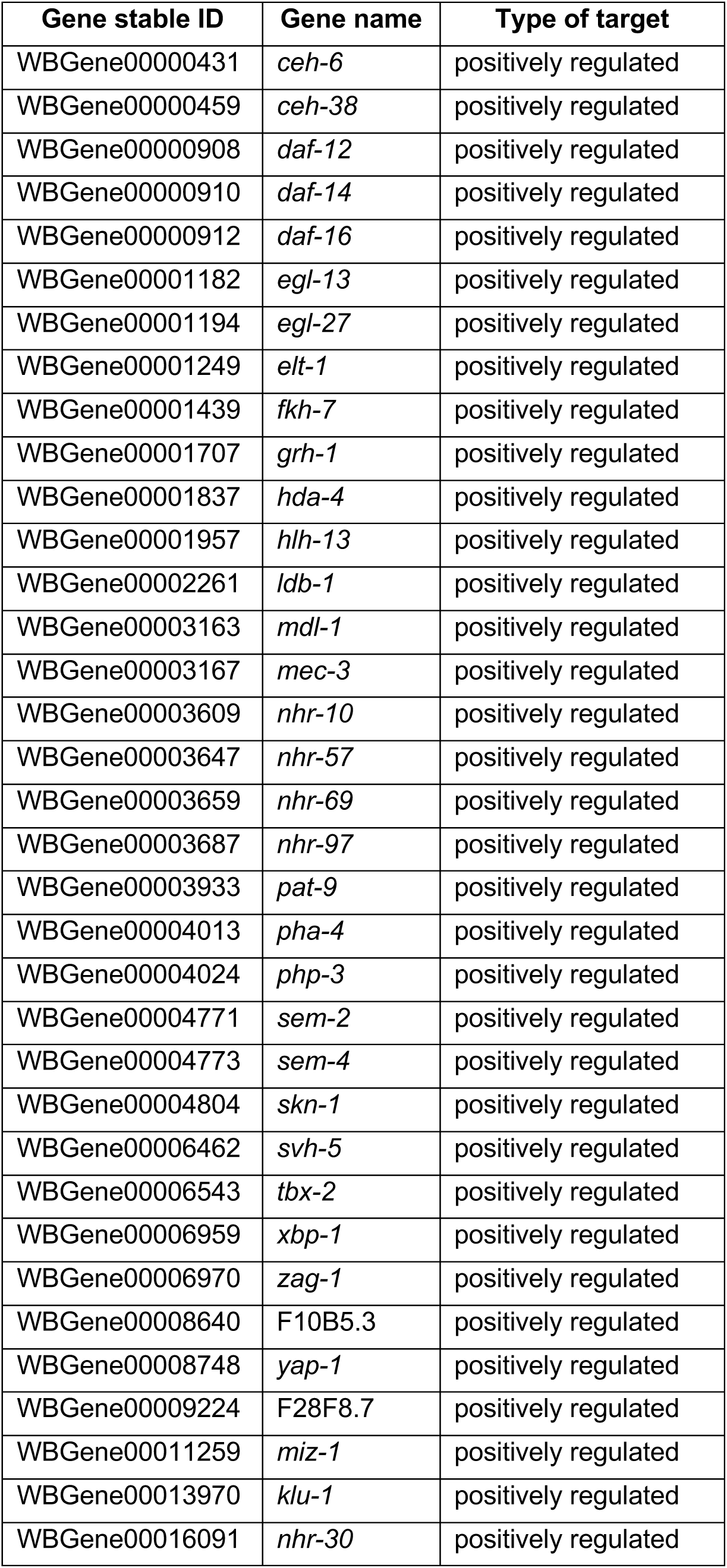
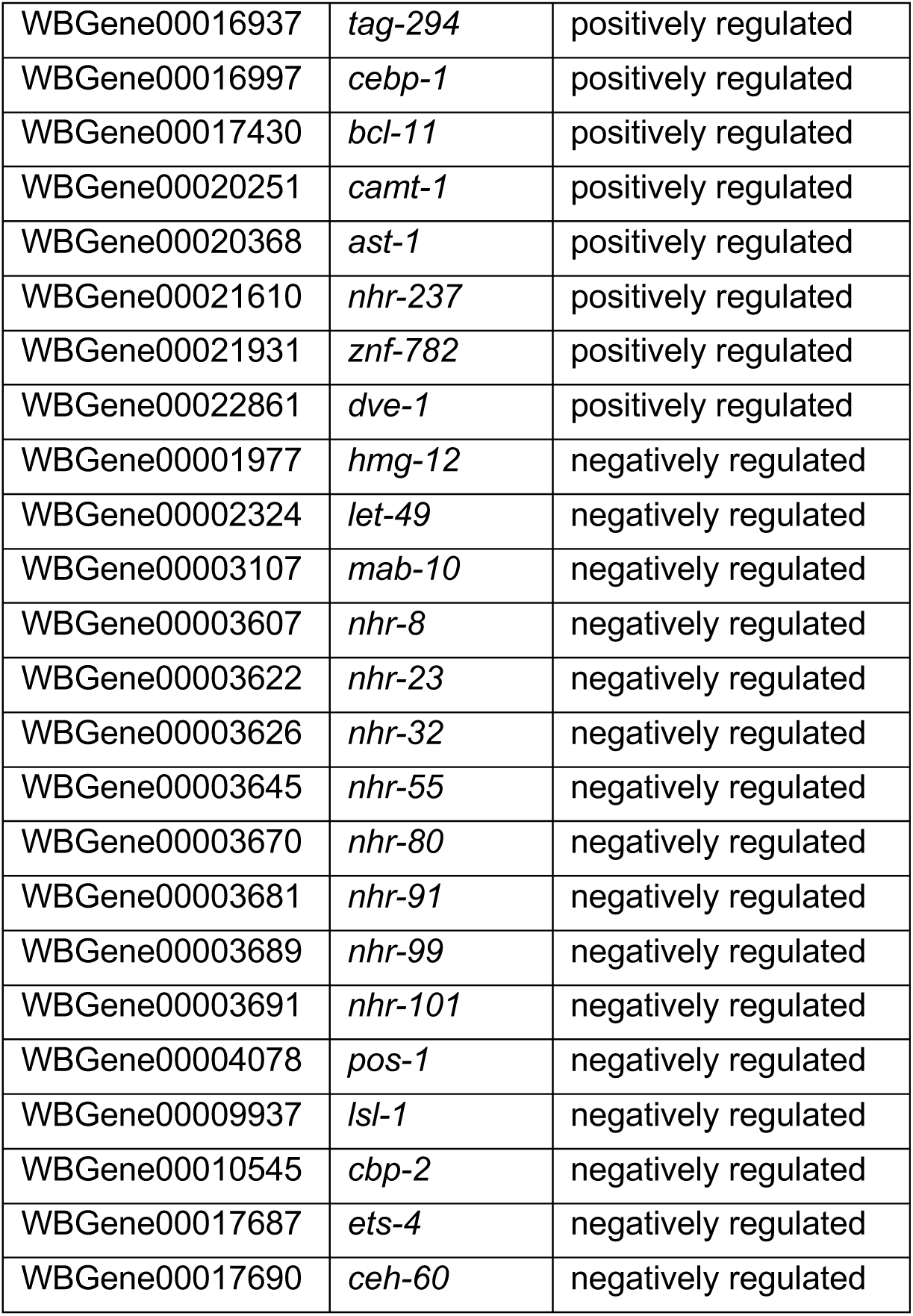
Transcription factors regulated by DAF-16.

**Table S2.** List of strains used in this study.

| Strain name | Genotype | Used in figures | Source/Reference |
| --- | --- | --- | --- |
| VT1777 | *daf-7(e1372) III; mals105[col-19p::gfp] V* | 1B, 1F, 1G, 2D, S1A, S2A, S2B, S4E, S6H | e1372: Riddle et al, Nature, 1981<br>mals105: Feinbaum, Ambros, Dev Biol, 1999 |
| XV36 | *daf-16(mgDf50) I; daf-7(e1372) III; mals105[col-19p::gfp] V* | 1B, 1C, 1D, 2D, S2A, S2B, S4E, S6H | mgDf50: Ogg et al, Nature, 1997 |
| XV226 | *daf-16(mgDf50)I; daf-7(e1372) III; mals105[col-19p::gfp] V; let-7(n2853) X* | 1C, 1D | n2853: Reinhart et al, Nature, 2000 |
| XV162 | *daf-16(mgDf50); daf-7(e1372) III; nDf51 mals105[col-19p::gfp] V* | 1C, 1D | nDf51: Abbott et al, Dev Cell, 2005 |
| QK244 | *daf-7(e1372) III; mals105[col-19p::gfp] V; xkEx51[col-183p::let-7::let-7 3'UTR + myo-2p::mCherry::unc-54 3'UTR]* | 1F, 1G, S1A | xkEx51: this study. Look at Methods section for details |
| QK152 | *daf-7(e1372) III; xkls43[let-7p::gfp]* | 2A, 2B, S4A, S4B | xkls43: this study. Look at Methods section for details |
| XV285 | *daf-16(mgDf50) I; daf-7(e1372) III; xkls43[let-7p::gfp]* | 2A, 2B, S4A, S4B |  |
| QK221 | *nhr-23(wrd8[nhr-23::gfp::aid\*::3xflag]) I; ieSi57[eft-3p::tir1::mRuby::unc-54 3'UTR + Cbr-unc-119(+)] II; daf-7(e1372) III* | 2E, S2C | wrd8: Ragle JM, et al. Development. 2020<br>ieSi57: Zhang L, et al. Development. 2015 |
| QK222 | *nhr-23(wrd8[nhr-23::gfp::aid\*::3xflag]) daf-16(mgDf50) I; ieSi57[eft-3p::tir1::mRuby::unc-54 3'UTR + Cbr-unc-119(+)] II; daf-7(e1372) III* | 2E, S2C |  |
| QK216 | *nhr-23(xk40[nhr-23::aid::3xflag]) I; ieSi57[eft-3p::tir1::mRuby::unc-54 3'UTR + Cbr-unc-119(+)] II; daf-7(e1372) III; mals105[col-19p::gfp] V* | 2F, 4A, S3B, S3C, S3D, S3E, S4C, S4D | xk40: this study. Look at Methods section for details |
| QK215 | *daf-16(mgDf50) nhr-23(xk40[nhr-23::aid::3xflag]) I; ieSi57[eft-3p::tir1::mRuby::unc-54 3'UTR + Cbr-unc-119(+)] II; daf-7(e1372) III; mals105[col-19p::gfp] V* | 2F, 4A, S3B, S3C, S3D, S3E, S4C, S4D |  |
| QK218 | *daf-7(e1372) III; mals105[col-19p::gfp] V; let-7(xk45[scRORE1,2,3]) X* | 2G, 2H, S5I, S5J | xk45: this study. Look at Methods section for details |
| QK219 | *daf-16(mgDf50) I; daf-7(e1372) III; mals105[col-19p::gfp] V; let-7(xk45[scRORE1,2,3]) X* | 2G, 2H, S5I, S5J |  |
| GS8924 | *daf-16(ar620[daf-16::zf1-wrmScarlet-3xflag]) I* | 3A, 3B, 3C, 3D, 3E, S6A | Gift from Iva Greenwald |
| QK230 | *nhr-23(wrd33[nhr-23::30aa linker:mScarlet:SEC:3XMyC]) I; daf-7(e1372) III* | 3F | wrd33: Myles KM, et al. MicroPubl Biol 2023 |
| QK231 | *nhr-23(wrd33[nhr-23::30aa linker:mScarlet:SEC:3XMyC]) daf-16(mgDf50) I; daf-7(e1372) III* | 3F |  |
| QK232 | *daf-16(xk48[daf-16::3xflag(DBD/N215A R218A H219A S222A)]) nhr-23(wrd33[nhr-23::30aa linker:mScarlet:SEC:3XMyc]) I; daf-7(e1372) III* | 3F | xk48: this study. Look at Methods section for details |
| QK245 | *daf-7(e1372) III; mals105[col-19::gfp] V; xkEx52[col-183p::nhr-23::3xflag::nhr-23 3'UTR + myo-2p::mCherry::unc-54 3'UTR]* | 4B, S9B | xkEx52: this study. Look at Methods section for details |
| QK200 | *nhr-23(xk40[nhr-23::aid::3xflag]) I; ieSi57[eft-3p::tir1::mRuby::unc-54 3'UTR + Cbr-unc-119(+)] II* | S3A |  |
| XV288 | *nhr-23(xk40[nhr-23::aid::3xflag]) daf-16(mgDf50) I; ieSi57[eft-3p::tir1::mRuby::unc-54 3'UTR + Cbr-unc-119(+)] II; daf-7(e1372) III; xkIs43[let-7p::gfp]* | S4A, S4B |  |
| QK206 | *let-7(mg279) X; wls54[scm::GFP]* | S5B | mg279: Reinhart et al, Nature, 2000 |
| QK036 | *let-7(n2853) X; wls54[scm::GFP]* | S5B |  |
| QK246 | *let-7(xk45[scRORE-1,2,3]) X; wls54 line 1* | S5B | xk45: this study. Look at Methods section for details |
| QK247 | *let-7(xk58[scRORE-1,2,3]) X; wls54 line 2* | S5B | xk58: this study. Look at Methods section for details |
| JR672 | *wls54[scm::GFP]* | S5B |  |
| QK350 | *let-7(xk45[scRORE-1,2,3]) X* | S5C, S5D, S5E, S5F, S5G, S5H |  |
| XV229 | *daf-16(ar620[daf-16::mScarlet]) I; daf-7(e1372) III; mals105 V* | S6H |  |
| QK233 | *daf-16(xk48 [daf-16::3xflag(DBD/N215A R218A H219A S222A)]); daf-7(e1372) III; mals105[col-19p::gfp]* | S8B, S8C |  |
| QK234 | *daf-16(xk49 [daf-16::3xflag]); daf-7(e1372) III; mals105[col-19p::gfp]* | S8B, S8C |  |

**Table S3.**
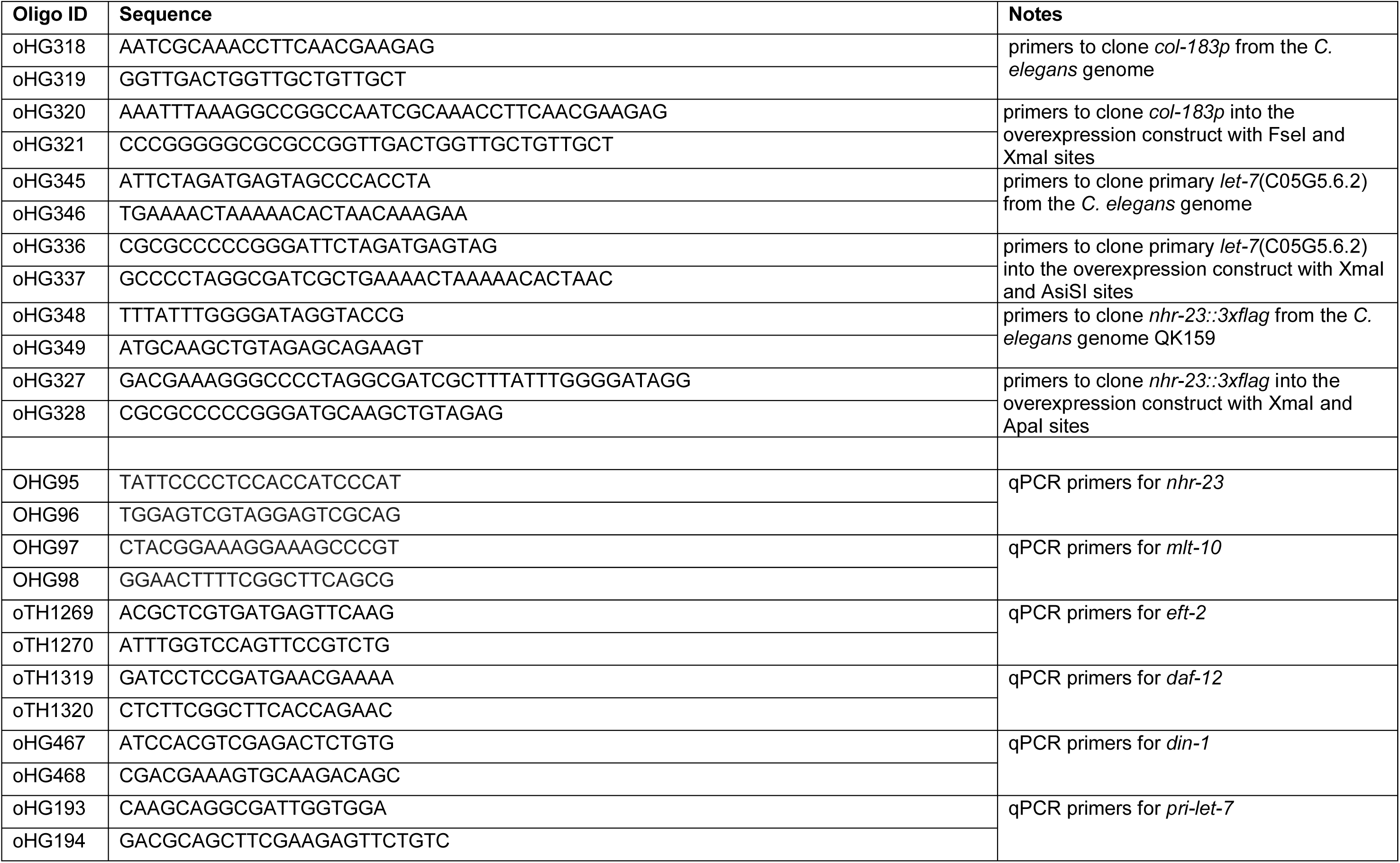
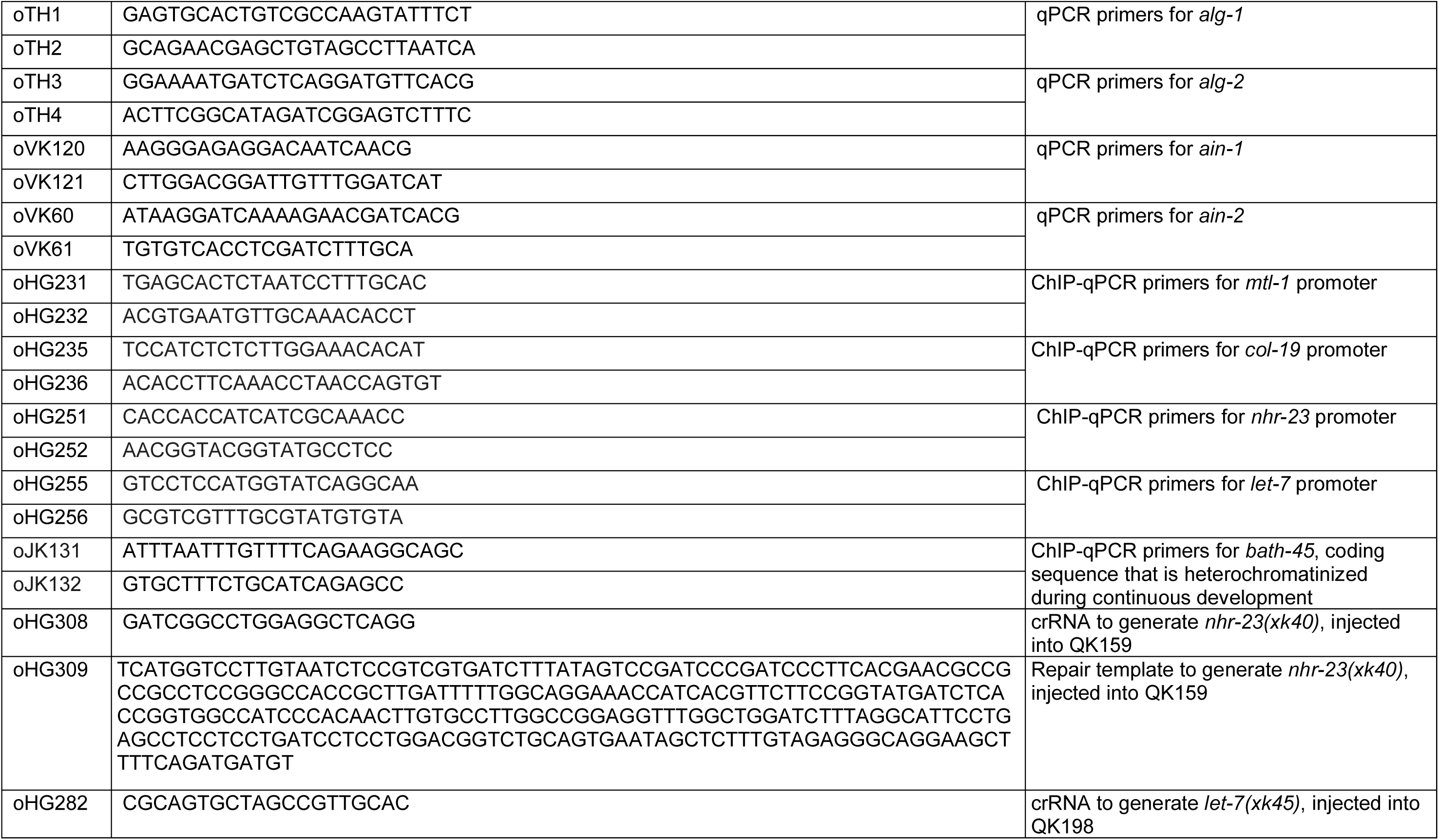
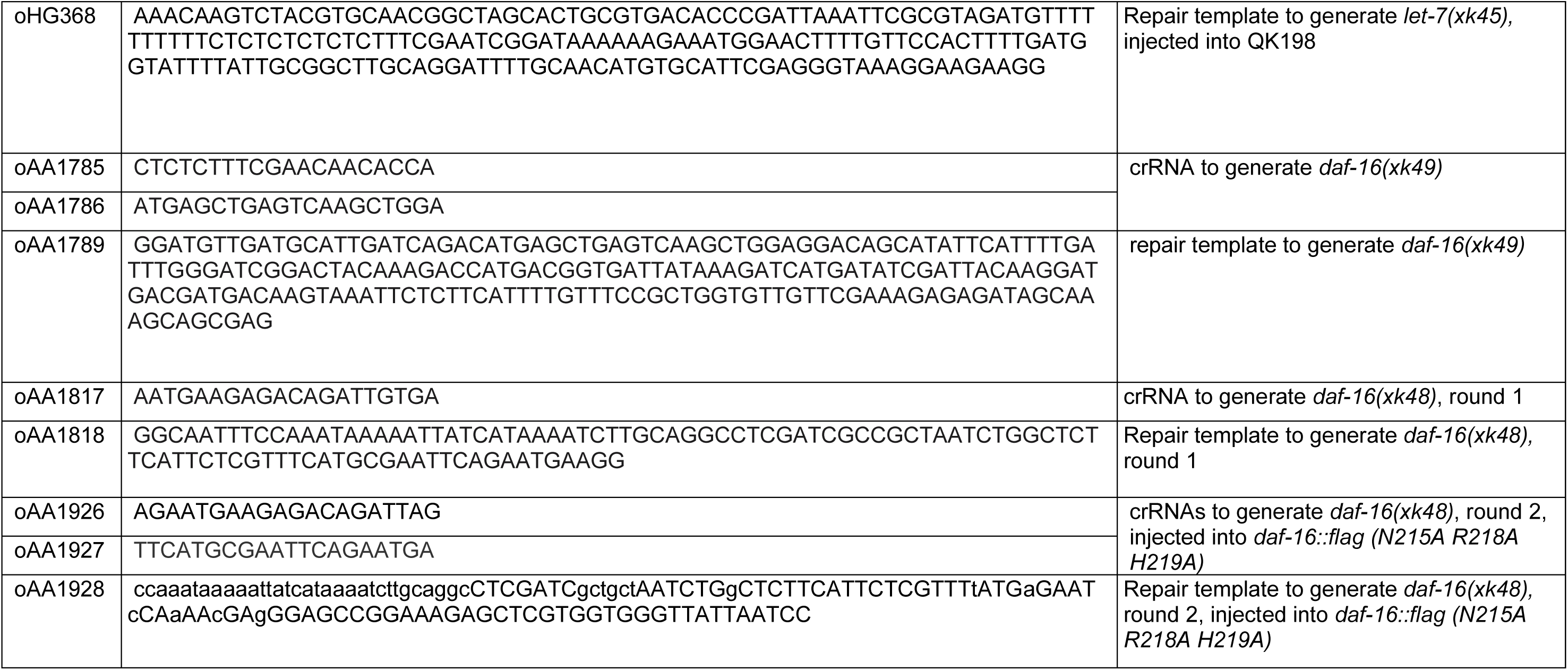
Oligonucleotides used in this study.

**Dataset S1 (separate file). Positively regulated targets of DAF-16 during dauer.**

**Dataset S2 (separate file). Negatively regulated targets of DAF-16 during dauer.**

**Dataset S3 (separate file). Gene Ontology analysis of the positively regulated targets of DAF-16.**

**Dataset S4 (separate file). Gene Ontology analysis of the negatively regulated targets of DAF-16.**

**Dataset S5 (separate file). RNA-seq analysis of Clock Controlled Genes and *nhr-23* targets.**

## Notes

https://www.ncbi.nlm.nih.gov/geo/query/acc.cgi?acc=GSE338973

https://www.ncbi.nlm.nih.gov/geo/query/acc.cgi?acc=GSE338292

### Competing Interest Statement

The authors have declared no competing interest.

